# Where and how our brain represents the temporal structure of observed action

**DOI:** 10.1101/276535

**Authors:** R.M. Thomas, T. De Sanctis, V. Gazzola, C. Keysers

## Abstract

Reacting faster to the behavior of others provides evolutionary advantages. Reacting to unpredictable events takes hundreds of milliseconds. Understanding where and how the brain represents what actions are likely to follow one another is therefore important. Everyday actions are predictable sequences of acts, yet neuroscientists focus on how brains responds to unexpected, individual motor acts. Using fMRI we show the brain encodes sequence-specific information in the motor system. Using EEG, we show visual responses were faster and smaller for predictable sequences that recruit the motor system. This study shifts the study of action observation from single acts to motor sequences, informs how we adapt to the actions of others and suggests the motor system may implement perceptual predictive coding.

## 1. Introduction

Perceiving and predicting the actions of other individuals is important to enable social interactions. Over the past decades, much work has been devoted to identify the neural mechanisms involved in processing simple acts such as grasping, reaching, breaking, and performing simple gestures. Electrophysiological work in monkeys has identified that mirror neurons in premotor and parietal regions involved in performing these simple acts are also activated when viewing or hearing others perform such acts (Gallese, Fadiga, Fogassi, & Rizzolatti, 1996; Umiltà, et al., 2001; Kohler, et al., 2002; Keysers, et al, 2003; Fogassi et al, 2005), and that the firing of individual mirror neurons contains information that permits accurate classification of which of two acts someone else is performing with high accuracy (C Keysers et al., 2003). Neuroimaging studies in humans have identified an *action observation network* triggered by the observation of such acts that involves visual regions of the occipital and temporal lobe on the one hand, and posterior parietal, premotor, cerebellar and somatosensory regions involved in performing similar actions on the other (Caspers et al., 2010; V. Gazzola and Keysers, 2009; Keysers and Gazzola, 2009; Rizzolatti and Sinigaglia, 2010a). This second set of regions has been referred to as the putative mirror neuron system (pMNS) due to its homologies with mirror regions in the macaque, with ‘putative’ acknowledging that we cannot know how much of the BOLD signals in this network directly stems from mirror neurons (Keysers and Gazzola, 2009). Using pattern classification analysis, some of these studies have confirmed that the pattern of brain activity in premotor, inferior parietal and somatosensory cortices indeed contains information about which motor act was performed by other (Etzel et al., 2008; Oosterhof et al., 2010).

In contrast, we know very little about **where** and **how** the brain represents **sequences** of acts (e.g. preparing breakfast Grafton and Hamilton 2007; Kilner and Frith 2008; Thioux et al. 2008), which inevitably will contain information that goes beyond the sum of those about each individual act. Representing sequences of acts means representing information about the order in which the acts were performed. Such sequence information is critical to predict what action people are likely to perform next, and hence, to start planning our reactions ahead of time. Here we will therefore explore first, where and second, how the brain represents such sequence-level information.

To explore **where** the brain encodes sequence level information, we will localize regions responding differently to acts in a logical sequence (e.g. grasping a bun, cutting the bun, buttering the bun) and in a random sequence. Some have argued that any such higher-level information is unlikely to be represented in the motor system, but is more likely to be localized in more cognitive brain regions such as the mentalizing network including the medial prefrontal, rTPJ and STS (Brass et al., 2007; Caramazza et al., 2014; Kilner and Frith, 2008). Others, including ourselves, in the light of monkey studies showing that mirror neurons in the motor system are sensitive to expectations about upcoming actions (Fogassi et al., 2005; Umiltà et al., 2001), suggest that the pMNS could represent sequence-level information. This is also in line with observations that premotor cortices can represent sequences of stimuli in other domains (Fiebach & Schubotz, 2006; Schubotz & von Cramon, 2001; Schubotz, et al, 2004) Indeed, we have argued, that because we can see our own actions unfold, Hebbian learning in the synapses mutually connecting our visual and motor systems would necessarily come to encode the transitional probabilities across individual motor acts, and hence enable our pMNS to represent sequence-level information and predictions in a predictive coding framework (Keysers and Gazzola, 2014).

A powerful approach, to investigate this question, has been exemplified in the domain of language. Lerner et al. (Lerner et al., 2011) took a story, and presented it to participants in its intact form, or after cutting at the edge between words, and randomizing the order of the words. If brain regions are only sensitive to word-level information, randomizing the order of the words in the story should not alter brain activity. If brain regions are sensitive to higher, sentence or paragraph level information, randomizing the order of the words should destroy that information and perturb brain activity. Brain activity was then analyzed using intersubject correlations (ISC) (Hasson et al., 2012). ISC maps information about a stimulus in the brain in a model free fashion based on a simple logic. If a voxel has no information about a stimulus, its activity reflects spontaneous activity and will not be correlated in time with that of other participants exposed to the same stimulus. If a voxel’s activity is determined by a stimulus, activity across witnesses of the stimulus will be similar, and the intersubject correlation will be significant. Hence, the higher the temporal correlation between subjects in a voxel, the more evidence we have for that voxel to contain information about the stimulus. By comparing ISC of the intact and scrambled sentences, the authors then identified brain regions that show evidence of additional information/correlation when sentence level information was preserved, thereby identifying regions that contain the equivalent of what we would call word-sequence information.

Here we adapted this approach to localize brain regions containing action sequence-level information. We recorded movies of familiar daily actions lasting approximately 1 minute (Table 1). We then measured brain activity using fMRI in 22 participants while they viewed intact movies that contain sequence- and act-level information. We also cut the movies at the transition between acts, and randomized the order of the acts, and measured brain activity while participants viewed these scrambled movies containing the same act-level information, but with perturbed sequence-level information (Figure 1). We then localized brain regions that had different levels of ISC for the intact and scrambled movies to identify regions showing sequence-level information. It is important to note that not finding a region in this contrast is not evidence that this region fails to encode sequence-level information. In addition to the usual limitations regarding negative findings, this is because ISC identifies activations occurring at the same location and time across participants, and hence focuses on stimulus-locked processes. If different participants encode the sequence of the overall actions (e.g. making breakfast) at different points along the sequence, this would evade the ISC analysis, and a region could then be involved in encoding this form of sequence-level information without showing increased ISC. We will therefore supplement ISC analyses with analyses exploring average activity levels across the sequences to shed light on less stimulus-locked processes. However, if a region shows more ISC for intact compared to scrambled sequences, we have evidence that this region contains sequence-level information.

**Table 1:** List of sequences used as stimuli with total duration in seconds and number of motor acts.

|  | **Action** | **Seconds** | **Acts** |
| --- | --- | --- | --- |
| 1 | Inflating and tying a balloon. | 51 | 27 |
| 2 | Making a paper boat. | 94 | 32 |
| 3 | Preparing bread with butter and jam. | 79 | 40 |
| 4 | Sewing a button. | 66 | 42 |
| 5 | Writing a gift card. | 83 | 39 |
| 6 | Rolling a cigarette. | 72 | 30 |
| 7 | Arranging flowers in a vase. | 82 | 39 |
| 8 | Framing a picture. | 112 | 39 |
| 9 | Cleaning spectacles. | 69 | 38 |
| 10 | Cleaning a laptop screen. | 46 | 28 |
| 11 | Sending a letter. | 42 | 34 |
| 12 | Replacing battery in a torch. | 51 | 27 |
| 13 | Applying nail polish. | 49 | 23 |
| 14 | Squeezing oranges. | 62 | 40 |
| 15 | Sharpening a pencil. | 83 | 44 |
| 16 | Replacing a pillow cover. | 44 | 35 |
| 17 | Removing nail polish. | 64 | 32 |
| 18 | Preparing a sandwich. | 77 | 27 |
| 19 | Toasting bread. | 65 | 30 |
| 20 | Folding a shirt. | 38 | 20 |

**Figure 1:**
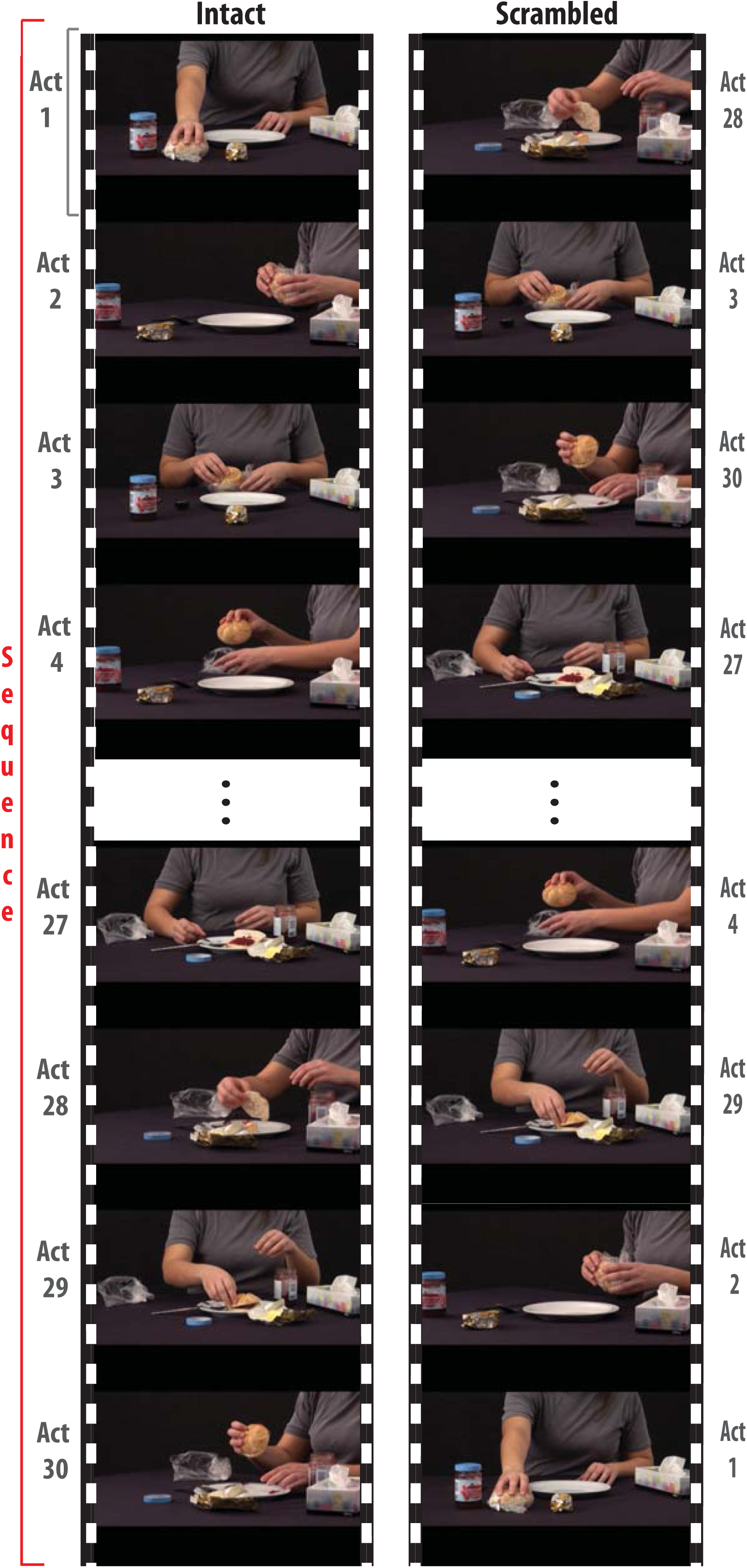
Stimulus used in the study. A movie of a familiar action (e.g. preparing a breakfast bun with butter and jam) is shown in an intact (left) and scrambled (right) version. Both versions contain the exact same individual acts (slicing the bun, spreading the jam, etc.), but in different order. Note the 45°camera angle change between every two consecutive acts in both intact and scrambled sequences. This was done to ensure that the inevitable visual transients created by rearranging a sequence in the scrambled condition are also present in the intact condition, and remove low level confounds.

Second, we aim to shed light on **how** the brain encodes sequence-level information. Anatomically, we know that areas in the higher levels of the visual system of the temporal lobe are reciprocally connected with regions of the posterior inferior parietal lobe which in turn are connected to dorsal and ventral premotor and somatosensory brain regions (Keysers and Gazzola, 2014; Keysers and Perrett, 2004; Nelissen et al., 2011). We can distinguish three families of models of the functional architecture of action observation based on how these models conceive of the feed-back connections back from parietal regions to the visual cortices (Figure 2). The first family emphasizes the role of feed-*forward* connections in triggering motor programs that match visual input (Rizzolatti and Sinigaglia, 2010b) without ascribing any specific function to the feed-back connections. The second family aims at explaining imitation and acknowledges the role of feed-*back* connections to visual regions, but assumes that they provide excitatory efference copies that *activate* matching visual representations in a way akin to mental imagery (Iacoboni et al., 2001). Finally, the third family, based on considerations derived from Hebbian learning and the observation that single neurons in the monkey STS are inhibited during action execution, proposes that neurons in the visual cortex are *inhibited* by parietal predictions via inhibitory feed-back connections (Keysers and Gazzola, 2014; Keysers and Perrett, 2004) in a way akin to predictive coding models derived from a Bayesian brain perspective (Kilner et al., 2007). Because in this last model, inhibitory feedback cancels predictions out of the visual response, the feed-forward visual information becomes a representation of prediction errors rather than of what is seen in the outside world. Here we will leverage the fact that these theories predict different activity patterns in the visual cortex to shed light on the computational mechanisms involved in action observation. Purely feed-forward accounts see the visual cortex merely as an input stage to action perception and therefore would not predict early visual areas to respond differentially to acts in and out of order. Excitatory efference-copy models would suggest that the response to individual predicted acts is amplified in early visual regions by excitatory efference copies, and hence that early visual responses should, if anything, be larger in intact compared to scrambled sequences. Inhibitory predictive coding models in contrast propose that early visual cortex essentially encodes prediction errors, and hence that early visual responses to individual acts should be largest in the scrambled sequences. In terms of the responses in the parietal node of the system, it is difficult to derive clear predictions from the first two families of models, but the predictive coding model predicts that the response to acts in intact sequence should be faster (by hundreds of milliseconds) than that of acts in scrambled sequences, because sensorimotor delays during re-afference wire the system so that preceding actions pre-activate expected sensorimotor representations in parietal cortex (Keysers and Gazzola, 2014; Keysers and Perrett, 2004). Because of its low temporal resolution fMRI is unsuited to resolve the individual motor acts embedded in our sequences, or sub-second shifts in response timing. Accordingly, to test these predictions we repeated our experiment using high-density EEG to compare the visual evoked response to individual motor acts in the intact and scrambled condition.

**Figure 2:**
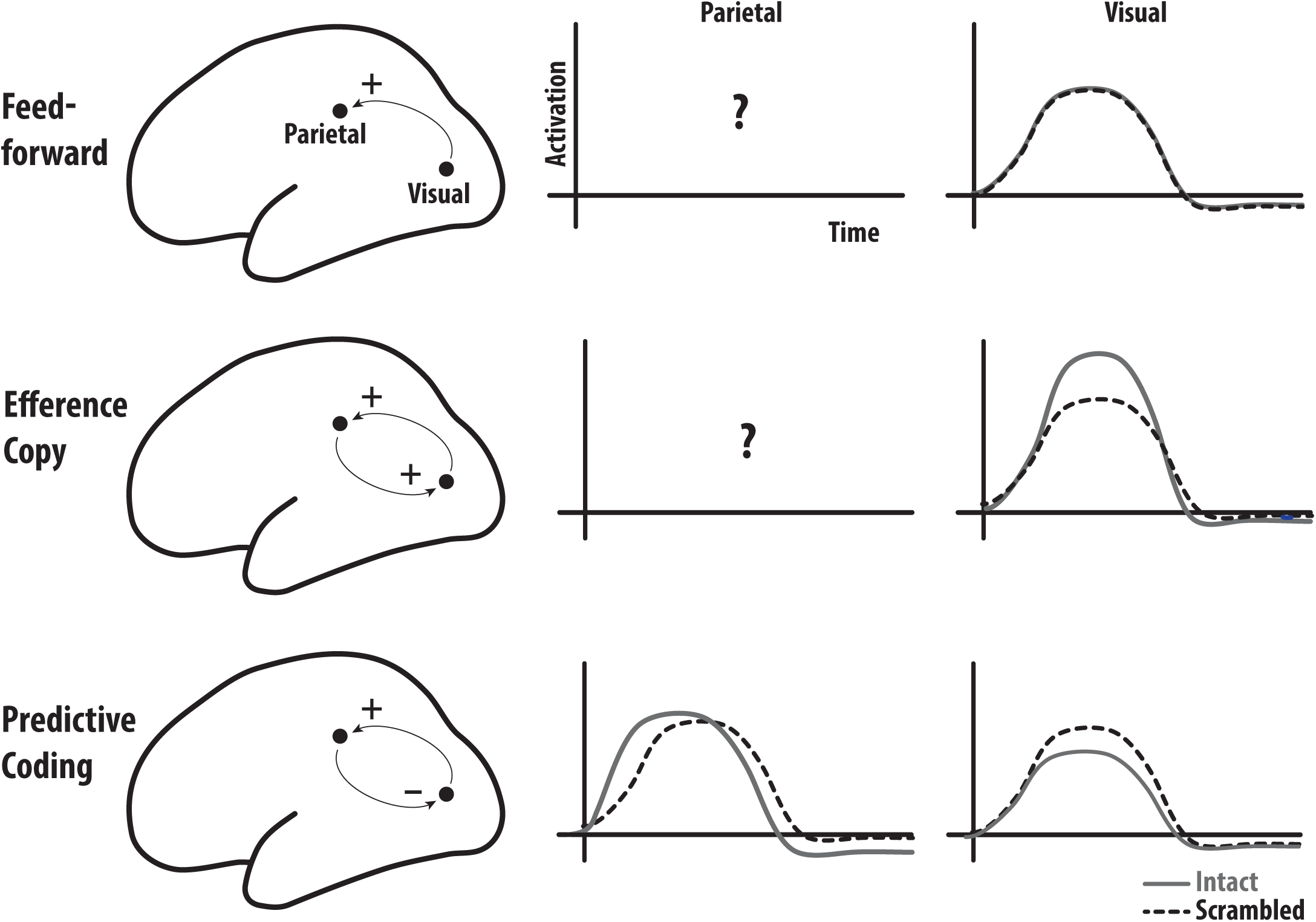
Predictions of different action observation models. Feed-forward models (top) emphasize feed-forward connections from visual to parietal regions and do not ascribe a function to feed-back connections. They do not make particular predictions on the timing of parietal activations for intact and scrambled sequences (middle column) but predict that visual cortices (right-most column) respond similarly to a particular observed act embedded in an intact and scrambled sequence. Efference-copy models (middle row) originating from imitation models suggest that feed-back connections are important and excitatory, and hence that in intact sequences, correct predictions in the parietal lobe should boost visual responses compared to scrambled sequences. However, it is unclear what predictions they make regarding the timing of responses in the parietal lobe. Finally, predictive coding theories suggest that in intact sequences, the parietal lobe should show predictive responses (that thus have latencies shorter than in scrambled sequences), that inhibit responses in the visual cortex (bottom row).

## 2. Results

### Intact movies show higher ISC

The ISC analyses revealed that both intact and scrambled movies (Figure 3 rows one and two, respectively) induce widespread synchronization across viewers. As might be expected, in both cases the visual cortices show high ISC reflecting the stimulus-locked nature of their responses. We also see significant ISC in parietal and premotor regions. The third row depicts the contrast between intact and scrambled (I-S). There were no significant voxels for which the scrambled movie shows a higher ISC than the intact movies. On the other hand, a number of areas show higher ISC for intact movies including the L/R postcentral gyrus (including BA2), L/R superior and inferior parietal lobule (including PF), and pre-central gyrus (including BA6) (Table 2).

**Table 2:**
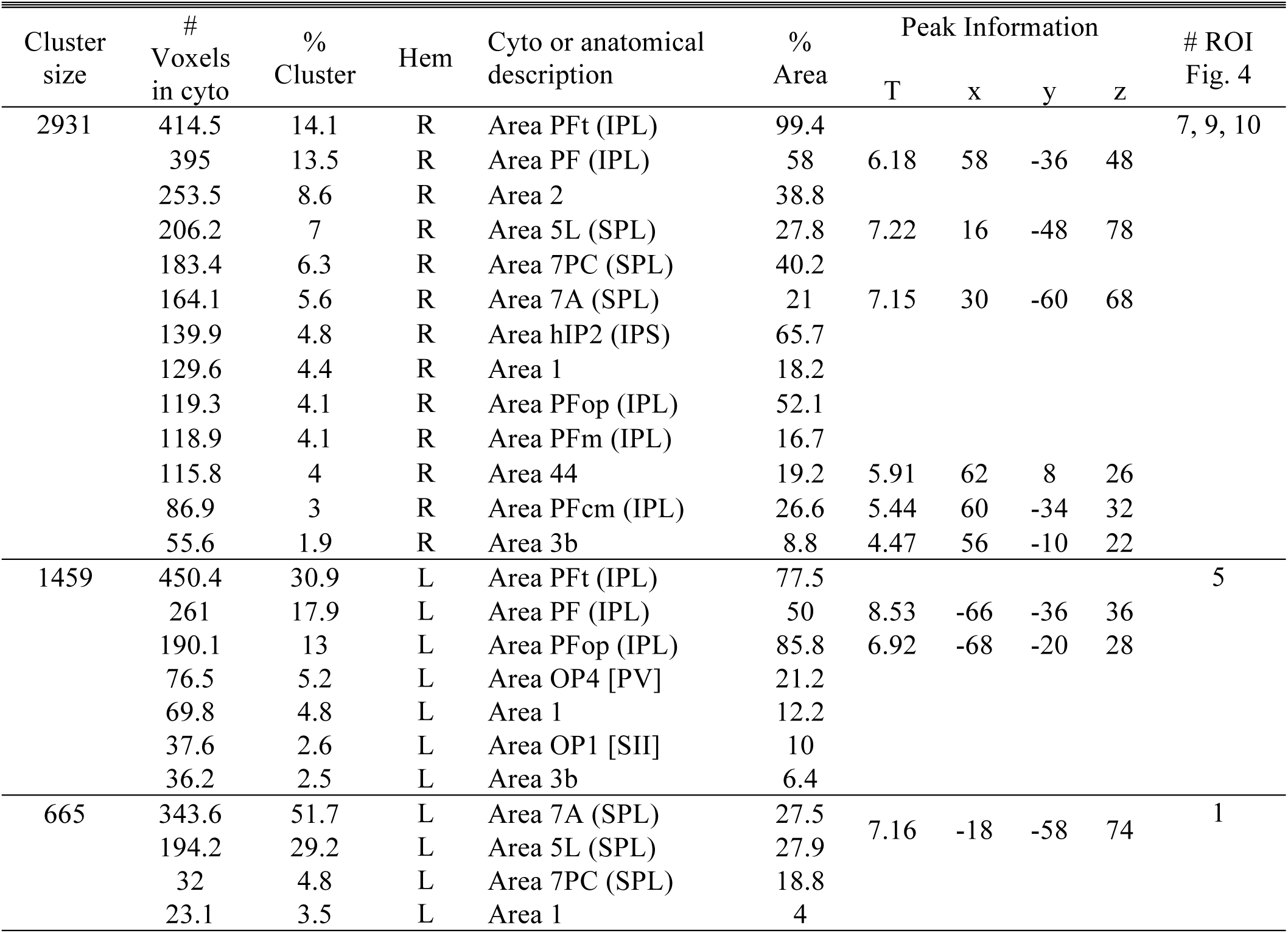
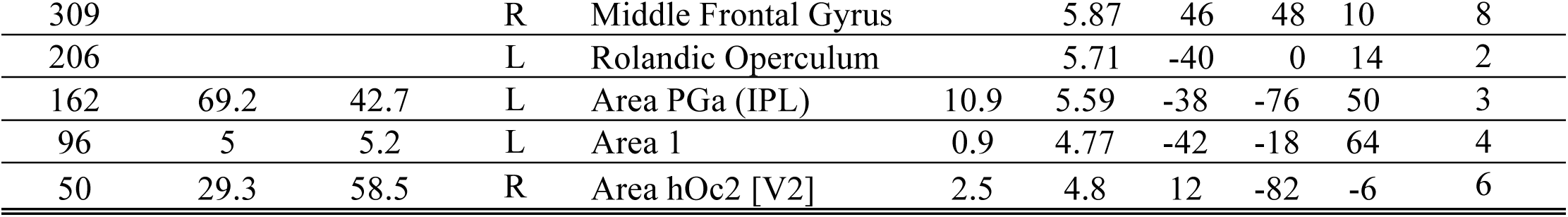
**ISC (I-S).** Regions with ISC Intact > Scrambled labeled using SPM Anatomy Toolbox. As for Figure 3, results are shown at p–values < 0.001 and cluster-size threshold of k=20 (to impose the same t>3.52 threshold on all contrasts), but voxels not surviving a voxel wise false discovery rate (FDR) correction at q = 0.05 were excluded (inclusive masking with in SPM). From left to right: the cluster size in number of voxels, the number of voxels falling in a cyto-architectonic area, the percentage of the cluster that falls in the cyto-architectonic area, the hemisphere (L=left; R=right), the name of the cyto-architectonic area when available or the anatomical description, the percentage of the area that is activated by the cluster, the t values of the peak associated with the cluster followed by the MNI coordinates in mm, the number of the ROIs in Figure 4 to which the cluster corresponds (the first cluster was split in 3 by increasing the threshold to t>4.1 to create these functionally more homogenous ROIs).

**Figure 3.**
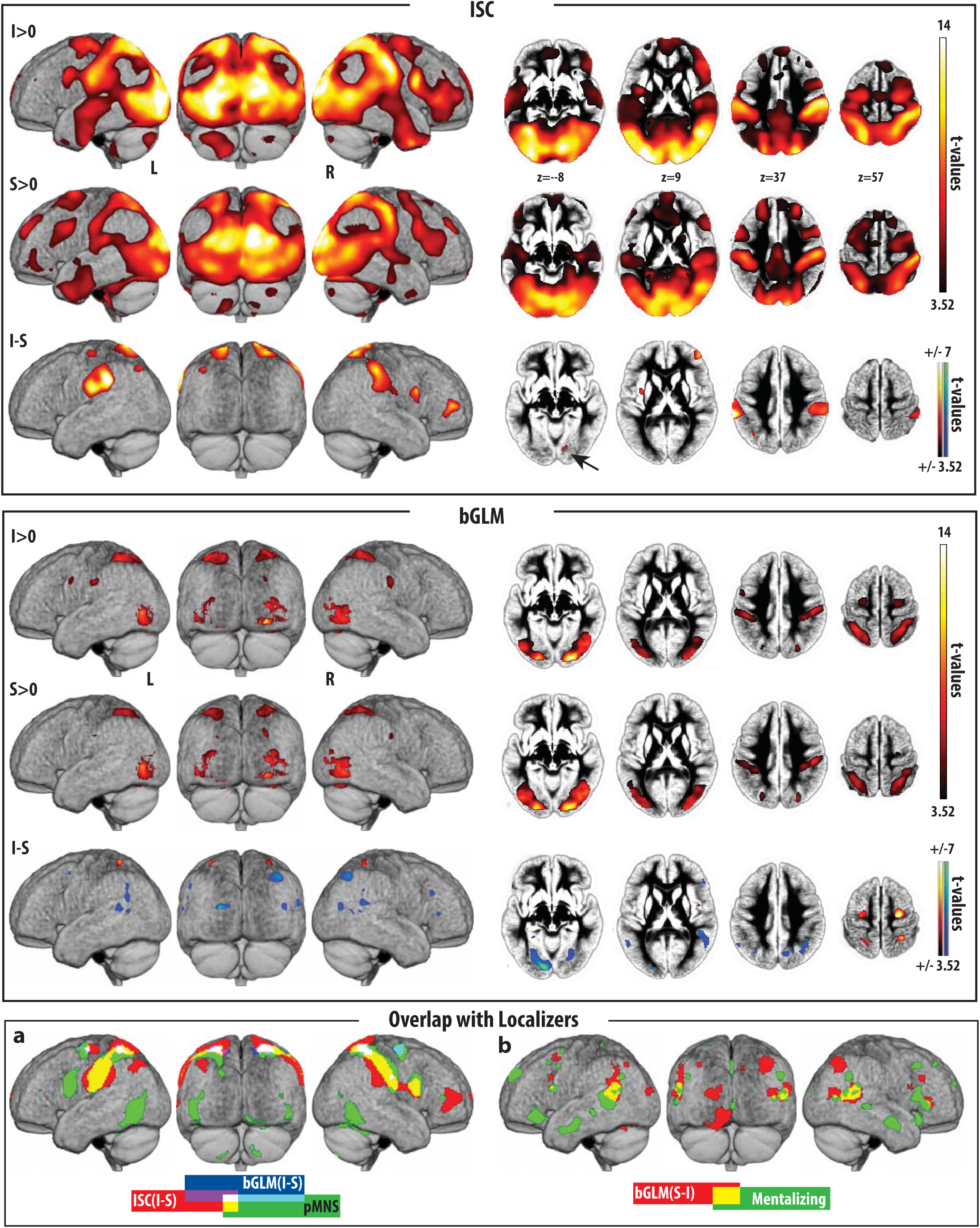
Regions with significant ISC (top panel) and bGLM (middle panel), and their overlap with the localizers. Each row of the top and middle panels corresponds to the contrast indicated on the left, and shows lateral and posterior renders and four axial slices on the average normalized gray matter segment of the 22 participants. Cold/warm colors represent significant negative/positive t-values. Results are shown at p–values < 0.001 and cluster-size threshold of k=20 (to impose the same t>3.52 threshold on all contrasts), but voxels not surviving a voxel wise false discovery rate (FDR) correction at q = 0.05 were excluded (inclusive masking with in SPM). The bottom panel shows the results of the overlap between the ISC and bGLM analyses, and our pMNS and mentalizing networks. (a) Overlap across regions showing more synchrony (ISC, red) or average activation (bGLM, blue) for intact movies and the pMNS (green). No overlap was found with the mentalizing network, which is why this network is not shown here. (b) Overlap (yellow) between regions showing more average activation (bGLM, red) during the scrambled movies and the mentalizing network (green), particularly in the TPJ. No overlap was found with the pMNS, which is why this network is not shown here. See Tables S1-S4, and Tables 2 and 4-6 for the corresponding MNI coordinate tables.

### Overlap between ISC (I-S), pMNS and the Mentalizing network

To investigate the degree to which I-S overlaps with regions associated with the putative mirror neuron system (pMNS), we used a functional localizer scan acquired in a separate group of participants (see Supplementary Methods S1). Briefly, it includes all voxels that are activated both when participants’ viewed goal directed acts (more than meaningless hand movements), and executed motor acts. The overlapping regions include right premotor (BA6), left and right inferior parietal regions (PF), and left somatosensory cortices (BA2) (Table 3; Figure 2 bottom panel). We also calculated the percentage overlap between pMNS and ISC (I-S) to find that 25.8% of all pMNS voxels show more ISC during I than S. To investigate the degree of overlap with the mentalizing network, we used the activation likelihood estimate meta-analysis of (Mar, 2011) that identified regions recruited by mentalizing based on non-story based tasks. No ISC (I-S) voxels were found to overlap with these mentalizing areas.

**Table 3:** **ISC (I-S) & pMNS.** Regions corresponding to the overlap between ISC (*I-S*) and the pMNS localizer, labeled using the SPM Anatomy Toolbox. Conventions as in Table 2. ISC data are thresholded as in Figure 3 and Table 2.

| Cluster size | # Voxels in cito areas | % Cluster | Hem | Cito or anatomical description | % Area |
| --- | --- | --- | --- | --- | --- |
| 943 | 366.8 | 38.9 | R | IPC (PFt) | 84.5 |
|  | 239.6 | 25.4 | R | Area 2 | 25.2 |
|  | 107.9 | 11.4 | R | IPC (PFop) | 40.1 |
|  | 70.4 | 7.5 | R | IPC (PF) | 7.9 |
|  | 65.6 | 7 | R | hIP2 | 45.9 |
|  | 15.5 | 1.6 | R | SPL (7PC) | 3.9 |
|  | 15.2 | 1.6 | R | Area 3b | 1.7 |
|  | 9.4 | 1 | R | Area 1 | 1.1 |
|  | 6.8 | 0.7 | R | OP 4 | 1.4 |
|  | 6.2 | 0.7 | R | hIP3 | 2.1 |
|  | 5.4 | 0.6 | R | OP 1 | 1.1 |
|  | 3.4 | 0.4 | R | IPC (PFm) | 0.4 |
| 490 | 169.4 | 34.6 | L | IPC (PFop) | 60.2 |
|  | 144.2 | 29.4 | L | IPC (PFt) | 35.9 |
|  | 109.6 | 22.4 | L | IPC (PF) | 11.2 |
|  | 37.2 | 7.6 | L | OP 1 | 6.3 |
|  | 12.5 | 2.6 | L | Area 2 | 1.4 |
|  | 8.6 | 1.8 | L | IPC (PFcm) | 2.2 |
|  | 6 | 1.2 | L | hIP2 | 2.5 |
| 153 | 64 | 41.8 | R | Area 44 | 7.2 |
| 48 | 21.6 | 45.1 | R | SPL (7PC) | 5.4 |
|  | 16.4 | 34.1 | R | SPL (7A) | 1.4 |
|  | 6.2 | 13 | R | Area 2 | 0.7 |
|  | 2.8 | 5.7 | R | SPL (5L) | 0.4 |
| 37 | 36.4 | 98.3 | L | SPL (7A) | 2.1 |

**Table 4:** **bGLM.** Regions with bGLM Intact > Scrambled, and Scrambled > Intact, labeled using SPM Anatomy Toolbox (Puncr<0.001, k>20 voxels, corresponding to t-values > 3.52). Conventions as in Table 2.

| Cluster size | # Voxels in cito | % Cluster | Hem | Cito or anatomical description | % Area | Peak Information |  |  |  |
| --- | --- | --- | --- | --- | --- | --- | --- | --- | --- |
|  |  |  |  |  |  | T | x | y | z |
| bGLM (I-S) |  |  |  |  |  |  |  |  |  |
| 430 | 323.2 | 60 | R | Area 6 | 7.4 | 7.81 | 26 | -14 | 58 |
|  | 180.4 | 41.9 | R | Area 2 | 27.7 | 4.23 | 38 | -34 | 44 |
|  | 151.6 | 35.3 | R | Area 7PC (SPL) | 33.3 | 6.33 | 30 | -46 | 58 |
|  | 54.1 | 12.6 | R | Area 5L (SPL) | 7.4 | 5.11 | 18 | -56 | 60 |
|  | 11.8 | 2.7 | R | Area 7A (SPL) | 1.5 |  |  |  |  |
| 340 | 130.4 | 38.3 | L | Area 5L (SPL) | 18.8 | 5.6 | -22 | -50 | 60 |
|  | 51.2 | 15.1 | L | Area 2 | 9.7 | 5.13 | -36 | -38 | 48 |
|  | 48 | 14.1 | L | Area 7A (SPL) | 3.8 |  |  |  |  |
|  | 37.6 | 11.1 | L | Area 7PC (SPL) | 21.9 | 6.05 | -32 | -48 | 66 |
| 311 | 251.5 | 80.9 | L | Area 6 | 5.7 | 6.37 | -26 | -16 | 52 |
| 23 | 13 | 56.5 | R | Area PFt (IPL) | 3.1 | 4.19 | 56 | -18 | 32 |
| bGLM (S-I) |  |  |  |  |  |  |  |  |  |
| 1090 | 276.2 | 25.3 | L | Area hOc4v [V4(v)] | 38.3 | 6 | -24 | -76 | -16 |
|  | 265.1 | 24.3 | L | Area hOc3v [V3v] | 28.6 | 8.79 | -10 | -88 | -10 |
|  | 163 | 15 | L | Area FG1 | 62.6 | 4.78 | -30 | -68 | -18 |
|  | 112.9 | 10.4 | L | Area hOc1 [V1] | 5.6 |  |  |  |  |
|  | 91.1 | 8.4 | L | Lobule VI (Hem) | 4.8 | 4.36 | -34 | -64 | -22 |
|  | 67.8 | 6.2 | L | Lobule VIIa crusI (Hem) | 2.2 |  |  |  |  |
|  | 47.2 | 4.3 | L | Area hOc2 [V2] | 5 |  |  |  |  |
| 628 | 95.9 | 15.3 | R | Area PGa (IPL) | 12.9 | 5.12 | 64 | -48 | 20 |
|  | 51.1 | 8.1 | R | Area hOc4la | 5.8 | 3.78 | 54 | -68 | 12 |
|  | 29.6 | 4.7 | R | Area PGp (IPL) | 3 |  |  |  |  |
|  | 15.9 | 2.5 | R | Area PFm (IPL) | 2.3 |  |  |  |  |
| 431 | 58.1 | 13.5 | R | Area hIP1 (IPS) | 20.1 |  |  |  |  |
|  | 26 | 6 | R | Area hIP3 (IPS) | 5.7 |  |  |  |  |
|  | 24.4 | 5.7 | R | Area PGa (IPL) | 3.3 |  |  |  |  |
|  | 15.9 | 3.7 | R | Area PGp (IPL) | 1.6 |  |  |  |  |
| 420 | 145.1 | 34.6 | L | Area PGa (IPL) | 22.7 | 4.7 | -54 | -56 | 26 |
|  | 40.5 | 9.6 | L | Area PFm (IPL) | 7 | 4.55 | -56 | -58 | 40 |
| 174 | 96.8 | 55.6 | R | Area hOc4v [V4(v)] | 15.6 | 4.26 | 24 | -76 | -12 |
|  | 39.5 | 22.7 | R | Area FG1 | 15.6 | 4.7 | 26 | -64 | -8 |
|  | 32.4 | 18.6 | R | Area hOc3v [V3v] | 3.8 |  |  |  |  |
| 164 | 72.1 | 44 | L | Area hOc4d [V3A] | 12.6 | 7.53 | -20 | -92 | 14 |
|  | 34.9 | 21.3 | L | Area hOc3d [V3d] | 3.5 | 3.96 | -10 | -94 | 22 |
|  | 12.2 | 7.5 | L | Area hOc4lp | 1.4 |  |  |  |  |
| 99 | 1 | 1 | R | Lobule VI (Hem) | 0.1 |  |  |  |  |
| 81 | 27.2 | 33.6 | R | Area 45 | 2.6 | 4.4 | 52 | 32 | 8 |
| 70 | 35.9 | 51.2 | L | Area hIP3 (IPS) | 7.8 | 4.02 | -30 | -62 | 40 |
|  | 9.6 | 13.8 | L | Area PGa (IPL) | 1.5 | 3.78 | -34 | -68 | 46 |
| 64 | 2.1 | 3.3 | R | Area 44 | 0.4 |  |  |  |  |
| 37 | 1 | 2.7 | L | Area 44 | 0.1 | 3.81 | -42 | 10 | 20 |

### GLM reveals areas differentially more active during the scrambled movies

Because ISC focuses on stimulus-*locked* activity, to identify regions with differential stimulus-*induced* activity (e.g. triggered by the stimuli but at different times in different participants), we analyzed the data using an approach in which each movie was modeled as a block in a GLM (bGLM) to capture the overall average activity during the viewing of a movie. Results of the bGLM are shown in Figure 3 (middle panel) for I>0, S>0 and I-S. Note that for I-S (Table 4), we have both positive and negative t-values. Regions with higher average activity for I than S (warm colors) included somatosensory cortices (left and right BA2) and supramarginal parietal regions (right PF) with 54% of these voxels falling within the regions identified by the ISC(I-S) results. Regions with higher activity for Scrambled movies (cold colors in Figure 3 middle panel; Table 4) included the visual cortices (V1, V2, V3 and V4) and the right angular gyrus (area PG).

### Overlap between bGLM, pMNS and mentalizing circuit

As for the ISC, the bGLM contrast identifying preferential activation for intact sequences (I-S) revealed no overlap with the mentalizing network but did overlap (as the ISC I-S did) substantially (1438 voxels) with the pMNS in premotor (BA6), rostral posterior parietal (PF) and somatosensory (BA2) regions (Table5, Figure 2 bottom panel).

Regions showing preferential activation for scrambled sequences (bGLM S-I) overlapped in 170 voxels with the mentalizing network (left and right angular gyrus, i.e. area PGa, and right BA45) and minimally (34 voxels) with the pMNS localizer in visual brain regions (Fusiform Gyrus) (Table 5,6, Figure 2 bottom panel)

**Table 5:** **bGLM & pMNS**. Regions corresponding to the overlap between bGLM activation maps and the pMNS localizer, labeled using the SPM Anatomy Toolbox. Conventions as in Table 2

| Cluster size | # Voxels in cito areas | % Cluster | Hem | Cito or anatomical description | % Area |
| --- | --- | --- | --- | --- | --- |
| **bGLM (I-S) & pMNS** |  |  |  |  |  |
| 442 | 231.9 | 52.5 | R | Area 6 | 5.3 |
| 382 | 180 | 47.1 | R | Area 2 | 27.6 |
| 264 | 102.4 | 38.8 | L | Area 5L (SPL) | 14.8 |
| 258 | 204.5 | 79.3 | L | Area 6 | 4.6 |
| 23 | 13 | 56.5 | R | Area PFt (IPL) | 3.1 |
| 18 | 16.9 | 93.8 | R | Area PFop (IPL) | 7.4 |
| 14 | 9.5 | 67.9 | R | Area PFt (IPL) | 2.3 |
| 14 | 14 | 100 | L | Area PFt (IPL) | 2.4 |
| 12 | 0.4 | 3.1 | R | Area OP3 [VS] | 0.2 |
| 11 | 5.4 | 89.6 | L | Area 6 | 0.1 |
| **bGLM (S-I) & pMNS** |  |  |  |  |  |
| 20 | 17.9 | 89.4 | L | Area FG1 | 6.9 |
| 9 | 6.8 | 75.0 | R | Area hOc4la | 0.8 |
| 6 | 5.2 | 87.5 | R | Area FG1 | 2.1 |

**Table 6:** **bGLM (S-I) & mentalizing areas.** Regions corresponding to the overlap between bGLM (*S-I*) activation map and the mentalizing network, defined using SPM Anatomy Toolbox. Conventions as in Table 2.

| Cluster size | # Voxels in cito areas | % Cluster | Hem | Cito or anatomical description | % Area |
| --- | --- | --- | --- | --- | --- |
| 104 | 14 | 13.5 | R | IPC (PGa) | 1.4 |
| 62 | 13.5 | 21.8 | L | IPC (PGa) | 1.6 |
| 4 | 4 | 100 | R | Area 45 | 0.4 |

### Network interactions in ISC (I-S)

To further characterize the 10 brain regions showing sequence-level information (Figure 3 ISC (I-S), and Figure 4a), we explored the correlation across their time courses (after averaging the time courses across all participants to isolate stimulus-triggered activity). A k-means clustering revealed that they segregate into three main networks of activity (shown as red, green and blue in Figure 4b) based on their cross-correlation pattern. The “red” network includes bilateral supramarginal clusters (including BA2 and PF), right inferior frontal gyrus (BA45), right precentral gyrus (BA44) and left insula. The “blue” network comprise right visual cortices (including area V1, V2 and V3) and left angular and dorsal premotor cortices. The “green” network includes clusters in the bilateral superior parietal lobule (including 7A and 5L). Correlation between the Eigen-time courses of these three networks (Figure 4c) was positive between the red and green networks and negative between the other pairs.

**Figure 4:**
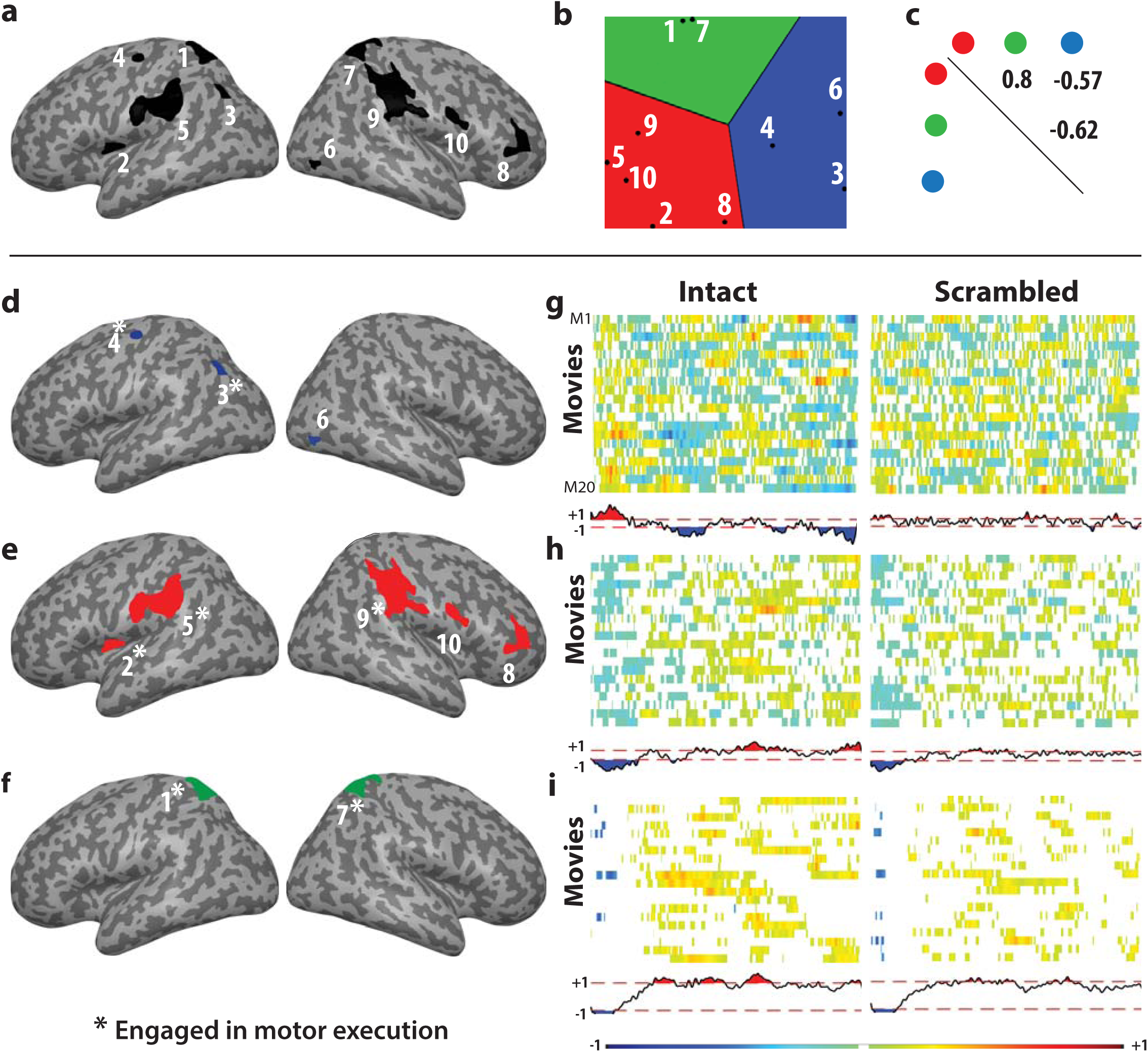
ISC Networks. The Eigen-time courses in the 10 regions showing ISC I-S (a) were compared using a *canonical multi-dimensional scaling algorithm* (b), in which regions with similar time courses are close to each other (see also Figure S1). A k-means clustering revealed that these regions can be summarized as 3 networks (colors in b). These networks (colored in red, green and blue separately in panels d-e) have significant correlations between them (c). (g-i) shows the average activity of these networks for each movie averaged over all participants (every row of the matrix corresponds to a movie) and the average across all movies (time course beneath each matrix). The white spaces in the matrix and the region bounded by red-dashed lines on the time course are where the ISC does not exceed that of randomized samples and is thus not significant. The ROI labels that have a star marked on top had significant co-activation with the motor-execution task (Supplementary Methods S2).

In order to have a better understanding of the kind of information encoded in these networks (Figure 4d-f), we plotted the Eigen time course of each of the network for each movie averaged across all subjects. By averaging the activity across all participants, one expects a flat line if activity is not synchronized across participants, and peaks (positive or negative) when the stimulus contains information that triggers activity in the same direction across participants. Figure 4g-i show a matrix for Intact (left) and Scrambled (right) that contains 20 rows, each corresponding to the brain activity triggered by a specific movie averaged across all subjects. To enable a more compact representation, the time course of activity is shown in a color code rather than an activation line, with time along the x-axis, and color intensity to signal the magnitude of activation, with warm colors signifying positive, and cold colors, negative activations. We further wanted to identify moments in which a movie activated a network beyond what is expected by chance. For this aim, rather than averaging the data of all the participants stemming from the same movie, we randomly permuted the labels of all the intact and scrambled movies, and averaged the activation again, repeating this procedure 1000 times. If a movie systematically triggered brain activity, one would then expect the original averaged activity to have a peak that is taller than those in the randomized movies. We therefore blanked all the parts of the matrix in which values remained within two standard deviations of the distribution of permuted data, while all the parts where values exceeded these bounds are shown in warm or cold colors.

Finally, to examine if there might be systematic trends across all movies, we performed a grand-average by averaging the activity across all movies, and showing them below the matrices, together with the bounds obtained from permuted grand-averages values (dotted lines). Note that the length of the movies has been rescaled to facilitate averaging across movies. This procedure was repeated for each of the three networks. In the case of the intact movies, finding moments of significant excursion within a movie is unsurprising, given that the regions of interest were selected to have strong ISC for the intact movies. That excursions remain significant after averaging across movies, however, indicates that the networks become synchronized consistently in specific moments of the movies. Interesting differences in timing between the three networks can be noted. The blue network, with its visual brain regions, is the first to show activation, while the other two networks are suppressed. This situation then reverses, with the activity of the green, and finally the red network becoming positive later in the movies. When processing the scrambled movies instead none of the networks show significant deflections except for the red network that shows negative ISC in the first part of the movie. We did not correct for multiple comparisons across time for this analysis, because the analysis aims to pinpoint when networks are most synchronized rather than establish that they synchronize, which would be circular given how ROIs were selected.

### Comparing ISC and bGLM results

We employed two streams of analysis (ISC and bGLM) to identify stimulus-locked and stimulus-induced activity, respectively. To better understand the relation between these method, we also extracted the parameter estimates of the bGLM from regions showing increased ISC, and inter-subject correlation values from regions showing altered bGLM (Figure 5, top row). In each case, we plotted the parameter estimates (beta values) for the ISC on the x- and the bGLM values on the y-axis. For each ROI, we then show the value for the intact movie as a circle, and for the scrambled movie as a cross. For ROIs selected based on increases in ISC, there is no point in statistically comparing the ISC value for intact and scrambled movies, as this would be circular. However, we tested whether there was a significant change in the bGLM, and showed those that were significant using a solid line (Figure 5 middle row), and those non-significant, with a dashed line. From this analysis, it becomes evident that parietal regions of the red network (ROI 4 and 10) increased their average brain activity in the intact compared to the scrambled movies along with the increase in ISC, while regions in the prefrontal cortex and visual regions (ROI 1, 6 and 8) reduced their average activity along with increases in ISC. The other ROIs did not change their average activity level at all, despite changes in ISC. For ROIs selected based on changes in bGLM, we tested the significance of the change in ISC. We see that most bGLM ROIs showing higher activity in the Intact condition (Figure 4, middle column) also showed higher ISC in the intact condition. In contrast, none of the bGLM ROIs showing more activity in the scrambled condition showed significant changes in the ISC (Figure 4, rightmost column). This suggests that activity triggered specifically by the intact movies, as identified using a GLM, was stimulus-locked (and hence leads to increased ISC), while that triggered specifically by the scrambled movies was stimulus-induced rather than -locked. Finally Figure 4 bottom row depicts the regions of overlap between ISC (I-S) with bGLM(I-S) and bGLM(S-I), respectively and shows that increases in ISC(I-S) can overlap with both bGLM increases (I-S) and decreases (S-I).

**Figure 5:**
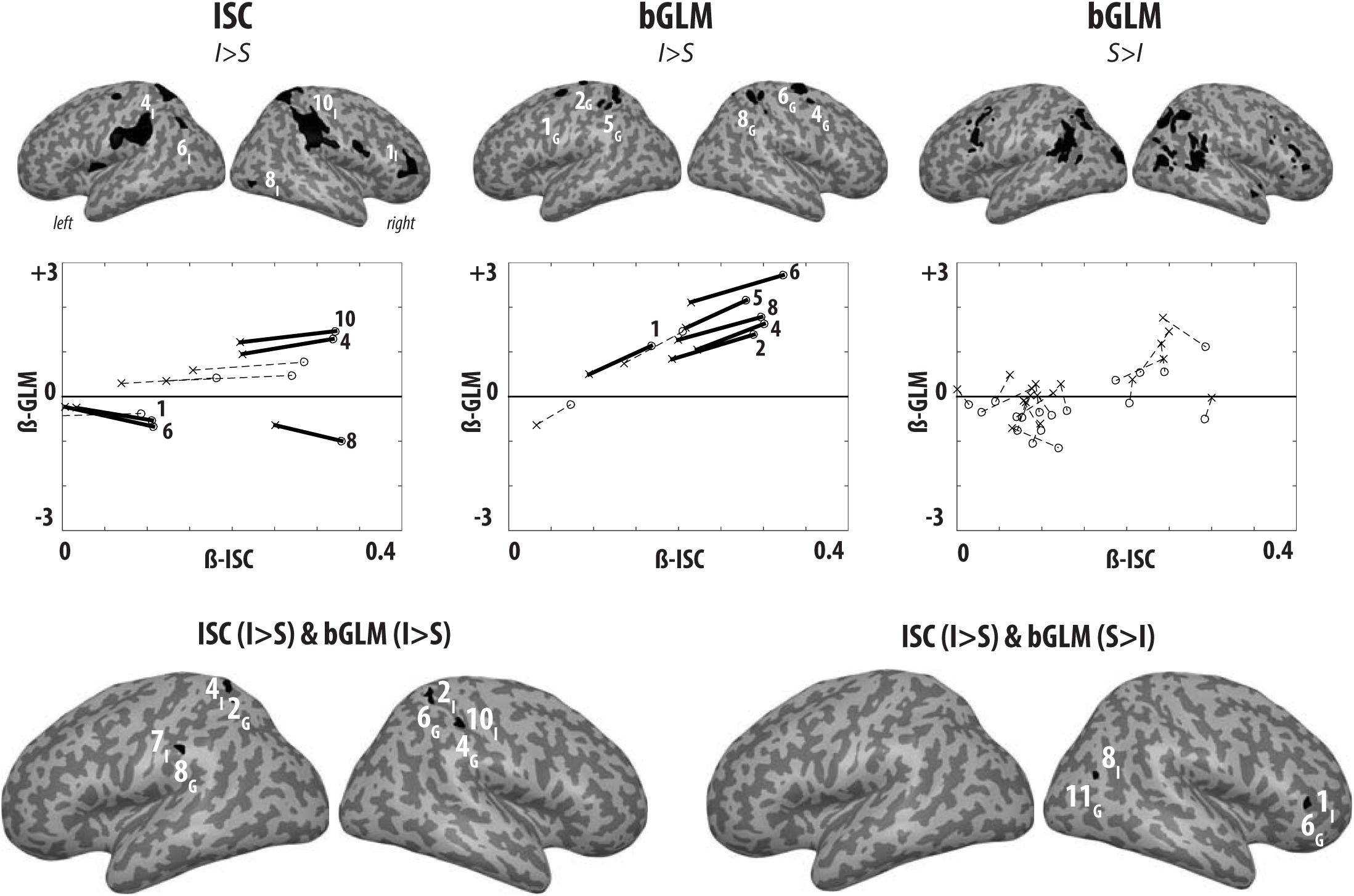
The ISC – bGLM relation. The top row depicts the ROIs corresponding to (from left to right) ISC, bGLM (I-S) and bGLM (S-I). The middle row the parameter estimates (*β* weights) from the ISC (x-axis) and bGLM (y-axis) as a function of the Intact/Scrambled state of the movie for the ROIs shown above. Significant differences between intact and scrambled in the dimension that was not used to define the ROI is shown by solid lines, non-significant differences, by dotted lines. The bottom row shows the overlap between ISC(I-S) with bGLM (I-S) and ISC(I-S) with bGLM (S-I). The numbers correspond to the ROI and the subscript I/G indicates membership to ISC/bGLM, so that 4I2G indicates an overlap between ROI 4 of the ISC and ROI 2 of the GLM.

### Intact movies lead to faster and smaller EEG responses compared to Scrambled movies

The different pMNS models make specific predictions regarding the responses to individual motor acts depending on whether they are embedded in intact or scrambled sequences (Figure 2). To test these predictions, we complemented the fMRI analyses with an analysis of electrophysiological responses triggered by each individual motor act, by aligning the EEG responses to the camera changes. Figure 6 shows the ERP that has been averaged across 5 distinct sets of EEG channels, progressively from posterior to anterior channels. We found that the ERP for intact movies started rising faster than for the scrambled movies, which was visible in the difference curves (Figure 6, insets) as an initial red phase. Additionally, we found that the ERP for intact movies had lower amplitude than that for the scrambled movies later in the ERP, which is visible as a blue phase in the difference curve.

**Figure 6:**
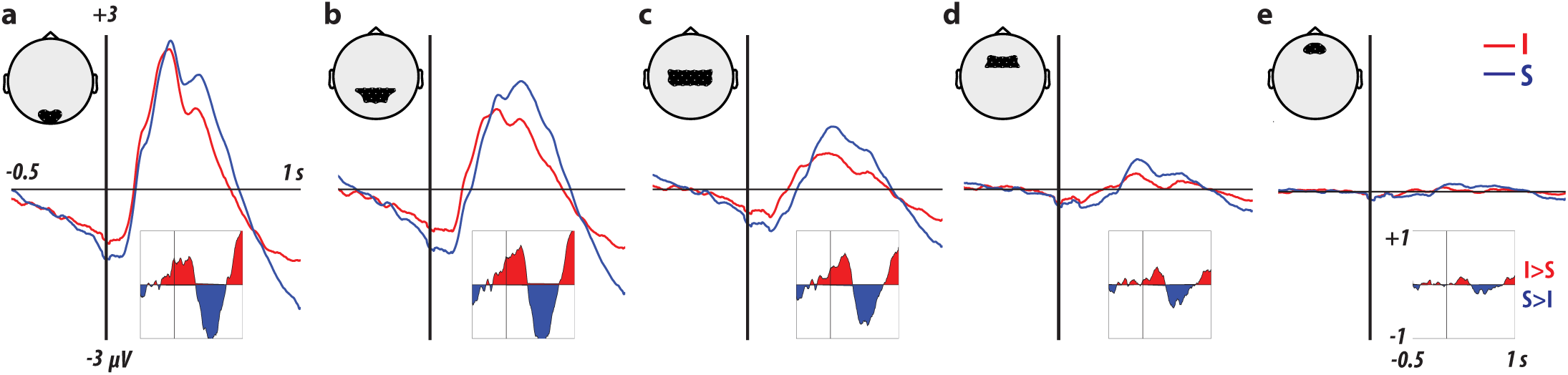
ERP for the intact (red) and scrambled (red) conditions. The inset below each ERP shows the difference curve (red for positive I-S and blue for negative) between intact and scrambled acts. The ERPs shown are also averaged across all the channels within the black shaded region shown on the sketch shown above each ERP. The channel groups enclose (a) OI1h, Oz, OI2h, O1, POO1, POO2, O2; (b) PO3, PPO1h, POz, PPO2h, PO4, P3, P1, Pz, P2, FP1, CPP5h, CPP3h, CPP1h, CPP2h, CPP4h, CPP6h; (c) CP3, CP1, CPz, CP2, CP4, CCP5h, CCP3h, CCP1h, CCP2h, CCP4h, CCP6h, C3, C1, Cz, C2, C4, FCC5h, FCC3h, FCC1h, FCC2h, FCC4h, FCC6h; (d) FC3, FC1, FCz, FC2, FC4, FFC3h, FFC1h, FFC2h, FFC4h, F3, F1, Fz, F2, F4; (e) AF3, AFF1h, AFF2h, AF4, AFP1, AFz, AFP2.

To quantify these observations, we performed two analyses. First, we explored the presence of a shift in latency and a difference in response magnitude using two simple models fitted separately to each participant’s ERPs. In Model 1, Scrambled(t)=Intact(t-λ), λ was optimized to reduce the residual error over the period from 0 to 300ms to focus on the rising of the ERP and quantify the shift in latency. In Model 2, Intact(t)= α *Scrambled(t), α was optimized to reduce residual errors for t from 0 to 650ms to take the entire response into account and quantify the relative response magnitude. We then compared the fitted parameters against the null hypotheses λ =0 and α =1 across participants using a cluster based statistic (see method). Figure 7a shows a significant shift in the response latency over occipital and parietal electrodes, with the intact condition leading to earlier responses (left panel) and a significant scaling over occipital, parietal and (left) frontal electrodes with the scrambled condition leading to larger responses. Second, we explored for time bins of 50ms, which electrodes show higher ERPs to intact than scrambled movies (Figure 7b) to confirm observations from the difference curves of Figure 6. Results confirmed that during the first 250ms, parietal and occipital electrodes show stronger responses to the intact movies, in line with the earlier rise time. From 250-400ms, no significant differences were observed. The pattern then reverses, with the scrambled condition triggering larger ERPs from 400-650ms over parietal and occipital electrodes. To provide an approximate mapping of the likely cortical sources for these differences at the electrode level, we performed a source localization of the ERP difference per time bin (Figure 7b right columns). The similarity of this source distribution over time suggests that a similar network of brain regions, including bilateral parietal and right visual cortices, is responsible for the earlier rise-time and reduced amplitude in the intact compared to the scrambled condition. (Given the limitations of source localization, we do not provide a coordinate table for these localizations).

**Figure 7:**
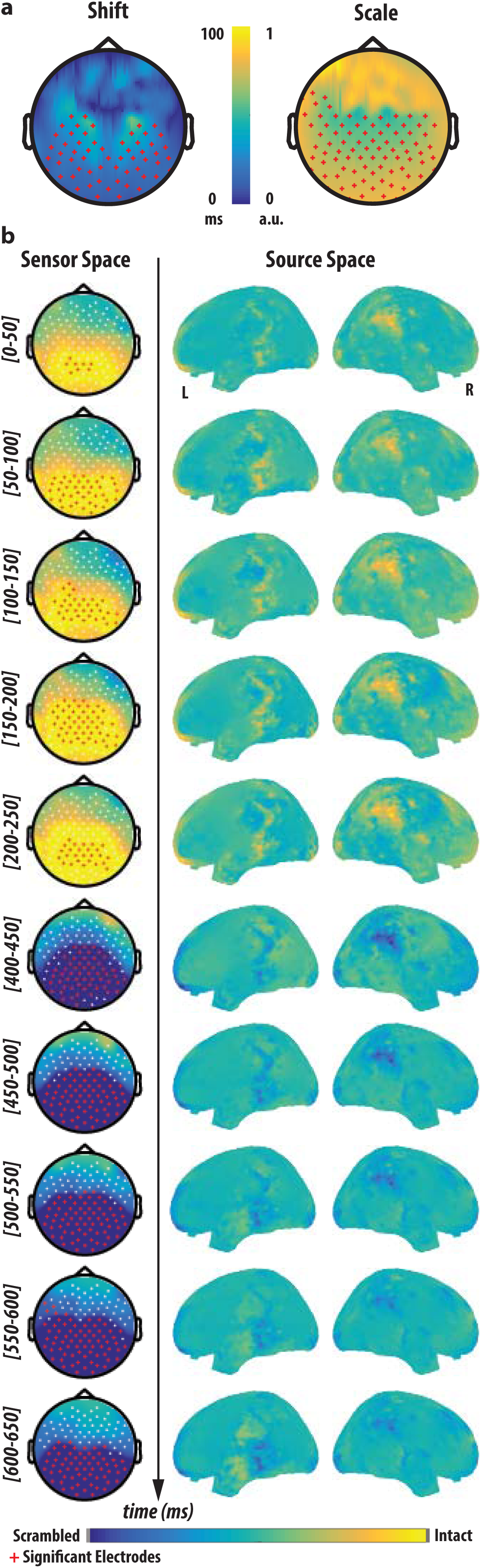
Differences in ERP across conditions. (a) *Topography of the parameters lamda (left) and alpha (right). Electrodes where lamda differs significantly from zero (with ERPs from Intact movies rising faster than those from Scrambled movies) or alpha differs significantly from one (with ERPs from Scrambled movies having higher magnitude) based on a cluster statistic are shown as red crosses. (b) Assessment of ERP differences as a function of time.* On the left the topoplot illustrates the difference in ERP between the intact and scrambled ERPs as a function of the time bin indicated. On the right side, a minimum norm source reconstruction of the difference. Blue indicates locations with Scrambled > Intact; yellow those were Intact > Scrambled. In all cases, sensors with significant differences between Scrambled and Intact (corrected for multiple comparison using a cluster statistic at p<0.05) are marked in red.

#### Discussion

In this experiment we set out to explore where and how the brain encodes sequence-level information when motor acts occur in sequences. Participants passively viewed a sequence of motor acts either in the natural (intact) or scrambled order while we measured their BOLD response using fMRI and their electrophysiological responses using EEG.

### Mapping sequence level information using fMRI

Regions that show higher ISC during the intact movie included three functional networks. One of these networks became active early in the movies (“blue” network in Figure 4d) and comprised right visual cortices (including area V1, V2 and V3) left angular (PG) and dorsal premotor cortices often jointly associated with stimulus driven spatial attention within the dorsal attention network (Nozawa et al., 2014, Nee et al., 2014). The other two networks became activated later when movies became more predictable and were negatively correlated over time with the first network. They included the “red” network in Figure 4e, formed by bilateral parietal clusters (including the PF complex in the rostral inferior parietal lobe and Area 2 and 1 in the primary somatosensory cortices) the left mid-dorsal insula and right inferior frontal (BA44) and middle frontal gyri, many of which are also involved in action execution (Gazzola & Keysers, 2009). They also included the “green” network composed of the superior parietal lobule bilaterally (including 7A and 5L), often associated with attention (Appelgren & Bengtsson, 2014, Chevrier et al., 2015). As mentioned in the introduction, there is debate in the literature about whether information beyond single motor acts should fall within the pMNS or the mentalizing network. We therefore explored whether these networks overlap with the pMNS (as localized in a separate group of participants) and/or with the mentalizing network (as localized from a meta-analysis).

### Overlap between sequence-level information and the pMNS

We found substantial overlap with regions included in our pMNS localizer. Regions of higher ISC during intact movies included in particular the classic pMNS nodes PF, the inferior frontal gyrus as well as the primary somatosensory cortex. Given that the intact and scrambled movies were composed of identical movie segments showing individual motor acts, if the pMNS were only to represent individual motor acts in isolation, we would expect the ISC to be identical in the two conditions. That the ISC was higher for the intact movies shows that the brain activity in these regions is not only sensitive to the individual motor acts, but also to their transitions, providing evidence that this network contains information at the sequence level beyond individual motor acts. This finding dovetails with predictions made by Hebbian learning models of the pMNS that suggest that this system should encode the transitions between actions (Keysers and Gazzola, 2014). It should be noted, that for brain activity triggered by a stimulus to translate into an increase in ISC, this brain activity must have two features: it must occur at the same location across individuals (after normalization and smoothing) and at approximately the same time across participants. The significant ISC increase we observe in the pMNS thus suggests that information about the transition between acts in the pMNS has – at least in part – these features. This is in line with the reliable timing observed in single mirror-neuron activity when observing single motor acts (C. Keysers et al., 2003; Kohler et al., 2002) and the consistency in the location activated by the sight of motor acts across individuals (Valeria Gazzola and Keysers, 2009).

That pMNS brain regions are sensitive to the sequence in which motor acts are observed supplements a small number of studies that show that other situational information that predict future acts can modulate mirror activity. Fogassi et al. have shown that if a monkey repeatedly witnesses someone grasping an object to bring it to the mouth, the activity of grasping mirror neurons is different from that in a context in which the monkey repeatedly sees someone grasp an object to place it in a container (Fogassi et al., 2005). Umilta et al. have shown that seeing a hand disappear behind an occluding screen that contains a graspable object leads to a higher discharge in grasping mirror neurons than seeing it disappear behind an occlude that doesn’t contain such an object(Umiltà et al., 2001). Finally, Iacoboni et al. showed that the ventral premotor node of the pMNS in humans responds differently to grasping a cup with a background of objects suggesting drinking vs. a background of objects suggesting washing the cup (Iacoboni et al., 2005). Together this suggests that the pMNS can take preceding acts, the objects in the scene and the situation into account in its response to a particular motor act, thereby endowing it with information that goes beyond single motor act. Further support for an association between the pMNS and sequence-level information stems from the fact that most of the regions that carried sequence-level information in our analysis were also activated when participants themselves manipulated objects (“*” in Figure 4) - a defining feature of mirror brain regions.

### Contributions of the mentalizing network

In contrast, we did not observe overlap between changes in ISC and the metalizing network. However, a bGLM analysis that compares the average activity during the entire movie in the intact and scrambled condition revealed that some voxels in the mentalizing network show higher average activation in the scrambled condition, but this did not translate into higher ISC in this network in that condition. Jointly this suggests that in contrast to the pMNS that becomes preferentially recruited in a stimulus-locked way when acts are embedded in natural sequences, the mentalizing network becomes preferentially recruited when motor acts deviate from natural sequences and then at times that vary across participant. This finding dovetails with the observation that mentalizing brain regions come online when witnessing implausible actions (Brass et al., 2007). Asking whether sequence-level information is represented in the pMNS or the mentalizing system is thus misleading. Rather these two systems become recruited by sequence-level information under different circumstances (Keysers and Gazzola, 2007).

### Evidence for inhibition of predictable visual responses

The second aim of this project was to contrast the predictions of different families of action observation models (Figure 2). Our EEG data supports inhibitory feedback models by showing that the amplitude and latency of the response to individual motor acts is reduced for intact compared to scrambled sequences. Source localization emphasizes the role of the parietal and visual cortices in generating these changes in timing and amplitude. Based on theoretical considerations alone, we had expected the changes in amplitude to be clearest over the visual cortex, and changes in latencies over parietal and premotor cortices – a separation that was not as evident in the EEG data where both changes were observed over occipital and parietal electrodes and the network identified in the source localization remained constant across time. This might reflect the technical limitations of EEG in separating the activity of individual nodes or suggest that in the sustained regime of activity induced by sequences of acts, the action observation network may start to act as one, more unified system rather than a series of individual nodes with distinct temporal properties – as is not unsurprising for a reciprocally connected loop. However, over the slower temporal scale of fMRI, the visual and parietal nodes of the pMNS had distinguishable time courses that lead them to fall within separable networks, with the parietal nodes peaking in activity later than the visual nodes. With regard to the distinction between inhibitory or excitatory feed-back between parietal and visual cortices, the observed anticorrelation between these fMRI networks is also compatible with the inhibitory-feedback model that would posit that the neural representation of observed actions should shift from visual brain regions initially to parietal and premotor regions that generate increasingly constrained and hence accurate predictions based on the somatosensory and motor connectivity pattern of the participants own actions in the intact sequences (Keysers and Gazzola, 2014). The bGLM also shows that a number of brain regions associated with visual processing show reduced BOLD signals while processing the intact sequences – a finding that is again most compatible with the notion of visual cortices receiving inhibitory feed-back in the context of intact sequences. The other models would not predict such negative correlations.

However, our data has limitations in its ability to establish predictive coding within the pMNS. First, our measurements cannot establish that the motor system is the cause for (i) the attenuation and acceleration of the responses in occipital electrodes in the EEG data for intact sequences or (ii) our ability to predict observed actions. Although it is compatible with such causal relationships, we need neuromodulation and/or lesion studies to establish such a causal relationship(Avenanti et al., 2017; Valchev et al., 2016). Second, a predictive coding framework posits that signals across visual and parietal cortices should provide *specific* predictions and prediction-errors about what action will come next, and how the observed action differs from these expectations – a specificity our data doesn’t establish. Designs in which we sequences recruit different body-parts (e.g. filling a glass then drinking it) in combination with pattern classification (Etzel et al., 2008; Oosterhof et al., 2010) might in the future address the specificity of these signals.

#### Conclusions

In summary our data suggests that when acts are arranged in sequences, the additional information is represented in separable networks based on whether the sequences adhere or not to the statistics of natural actions. When acts adhere to the natural statistics of our own actions, regions overlapping with the pMNS encode sequence-level information in a stimulus-locked fashion. An inhibitory feedback architecture, in which the visual processing of acts is inhibited by expectations derived from previous actions, is most compatible with the pattern of responses we observed: a reduction of visually evoked responses in EEG, a reduction in average BOLD activity in the visual cortex for intact compared to scrambled sequences and a negative correlation between BOLD activity in visual and pMNS nodes. When the order of the acts violates natural statistics, brain regions associated with mentalizing seem to encode sequence-level information in a spatially consistent but temporally more variable way, as seen in an increase of average BOLD activity without changes in ISC in this network. These findings invite brain activity manipulation and lesion studies that could investigate whether sequence processing would suffer more from perturbations of the pMNS when sequences fit natural statistics and mentalizing regions when they violate such statistics.

## 3. Materials and Methods

### Participants

All participants were right handed per the Edinburg Handedness Inventory (Oldfield 1971), had normal or corrected-to-normal vision and no history of neurological or psychiatric disorder. Informed consent was provided by each participant according to the procedure approved by the ethics review board of the University of Amsterdam. For the fMRI experiments, 22 healthy Caucasian participants took part (11 male, 11 female, mean age 23.3 ±3.46sd). None of the participants were excluded from the fMRI dataset. For the EEG experiment, a total of 24 participants were tested. Of these, 10 had also taken part in the EEG experiment (5 male). The other 12 fMRI participants could unfortunately not be traced back when we decided to perform the follow up EEG experiment, and an additional 14 participants were recruited for the EEG experiment alone (7 male, 7 female, mean 25.07±6.53sd). Three of these additional EEG participants were excluded due to excessive artefacts, leading to a final sample of 21 EEG participants (12 females, age: 26.76±5.86sd).

### Stimuli & Experimental Procedure

Twenty movies containing different daily actions (e.g. preparing bread with butter and jam; see Table 1 for the full list) were recorded simultaneously by two video cameras (Sony MC50, 29 frames/s) at an angle of 45°. The videos were edited using ADOBE Premier Pro CS5 running on Windows. Each movie was subdivided into shots containing one meaningful motor act each (e.g. taking bread, opening butter, scooping butter with knife, etc.). This was done on recordings from both camera angles. These motor acts (mean duration 2s ± standard deviation 1s) were then assembled to build two types of stimuli (Figure 1). For the so-called *intact* (I) movies, the natural temporal sequence in which the acts were recorded was maintained, but a camera angle change was introduced between every two consecutive acts by alternate sampling from the recordings of the two cameras. For the so-called *scrambled* (S) movies, the order of the acts was randomly rearranged, and a camera angle change also occurred between every two consecutive acts. Camera angle changes were imposed at each act transition in both types of movies to compensate for the visual transients that would otherwise be present only in the scrambled movies.

During both the fMRI and EEG experiments, participants had to carefully watch all the 20 movies, which were presented using the Presentation software (Neurobehavioral Systems, Inc., Albany, CA, USA) in four different sessions each containing 5 intact and 5 scrambled examples, shown in a pseudo-randomized fashion, with an inter-movie interval between 8 and 12 seconds. No behavioral response was required during the four sessions. To ensure that the participants were attentive to the movies and maintained a consistent engagement with them, they were asked to answer some simple questions regarding the stimuli being presented in the interval between two sessions (e.g., *Did you see roses or tulips during the movie clip?)*. They were not asked explicitly to pay attention to any particular detail.

### fMRI acquisition

Data were acquired on a 3T Philips Achieva scanner with a 32-channel head coil. Functional images were acquired with simultaneous multislice excitation equal to 3, (TR=721ms, TE= 28ms), 39 axial slices of 3mm with no gap and FOV of 240 × 240 × 39 mm. Images were reconstructed offline by Recon (Gyro Tools, Switzerland, http://www.gyrotools.com), after which the FOV was 120×78×240 mm. For each participant a T1 weighted image of 1×1×1 mm voxels was acquired. Stimuli were projected on an LCD screen and viewed through a mirror attached to the head coil.

### fMRI Inter-subject Correlation Analyses

Data were preprocessed using SPM12 (http://www.fil.ion.ucl.ac.uk/spm/software/spm12/) and custom-built MATLAB 9.8 (Mathworks Inc., Sherborn, MA) routines. The raw voxel time courses were bandpass filtered between 0.01 and 0.2 Hz, as this was shown to be the optimal band to perform ISC (Kauppi et al., 2010). At this stage the BOLD time courses corresponding to all the intact or scrambled movies presented to the subject were extracted, demeaned (voxel- and movie-wise) and concatenated such that, irrespective of the pseudo-random order in which the subjects saw the movies, the concatenated order was invariant across the participants. Before concatenation to a single 4D NIFTI file, we trimmed 3 TRs from the beginning and the end of each movie epoch to remove the influence of unspecific BOLD transients (Hasson et al., 2004). Each subject’s 4D file was then realigned to the mean image of the time course. The T1 weighted anatomical image was then co-registered to the mean functional image and segmented. All EPI images were normalized, at 2×2×2mm resolution, to the template MNI brain using the forward deformation tensor derived from the segmentation of the T1 image of that subject. The normalized images were then smoothed with an 8×8×8 mm (FWHM) Gaussian filter Inter-subject correlations were calculated using the ISC toolbox (Kauppi et al. 2014) and in-house MATLAB routines and SPM12.

After the preprocessing step for ISC we have two 3D time courses per subject, one for intact and the other for scrambled movies. For the subject-level ISC analysis, the time course of a given voxel in subject *i* was correlated with the average time course of all other subjects of that corresponding voxel. This was repeated for every voxel and in all subjects, resulting in a whole brain map of correlation values per subject (Hasson et al., 2010; Kauppi et al., 2014).

These correlation maps are then used in a second-level analysis in SPM: a one sample t-test for I>0 (H0: I<0) and S>0 (H0:S<0) and a paired sample t-test comparing I and S (H0:I-S) was performed at every voxel. Using a voxel-wise false discovery correction for multiple comparison with q<0.05 and cluster-size threshold set at k>20 voxels, t-values greater than 2.56 and 2.58 for I>0 and S>0 respectively and 3.92 for I vs. S are significant.

The ISC revealed that intact movies generated activity that was more synchronized across participants than scrambled movies in 10 clusters (Figure 3 ISC I-S). To determine whether these regions clustered in a smaller number of networks, we transformed these 8 clusters into 10 regions-of-interest (t>3.92 corresponding to q_FDR; k=20_ < 0.05, Figure 4a and Table 2) using MARSBAR (http://marsbar.sourceforge.net/) to extract the Eigen-time-course of each of them. Because the right parietal clusters was very large, and spanned several cyto-architectonic regions at q<0.05, we split this cluster in several sub-clusters by marginally increasing the threshold from t>3.92 to t>4.1, which split this large cluster into three clusters (rois 7,9 and 10 in Figure, 4). The Eigen time-course from each of these ROIs was then averaged over all participants to focus on stimulus driven activity and used to form a cross-correlation matrix between all the ROIs (see Supplementary Figure S1). This correlation matrix was used as input to a *canonical multi-dimensional scaling algorithm* that positioned the ROIs on a 2D map according to the pattern of correlations between a particular ROI and others, with ROIs having similar patterns mapped closer to each other. A k-means clustering algorithm was then used to determine the number and membership of clusters in this data. To determine the number of clusters, given the relatively low number of data-points, we used the Silhouette procedure (Kaufman and Rousseeuw, 1990). The procedure involves computing the k-means with k being set to 2,3,4 and 5 clusters (more clusters seemed inappropriate for 10 data-points). For each value of k we calculated the Silhouette value for each data point (large value implies the point is well within a cluster and low values reflects ambiguity).

### fMRI General Linear Model Analyses (GLM)

The preprocessing pipeline for the GLM analyses began with temporal filtering as in the case of the ISC. Subsequently the four different sessions were slice time corrected and realigned. The T1 image was co-registered to the mean EPI, and segmented. The EPI images were then normalized using the forward deformation tensors derived from that segmentation, written at a 2×2×2mm resolution, and then smoothed with an 8×8×8 mm (FWHM) Gaussian filter.

In the first level analysis of the GLM, the five blocks of the intact and five of the scrambled movies were modeled as two regressors-of-interest in each of the sessions. Movement parameters estimated during realignment were included as covariates of no interest in the analysis. The regression coefficients were then used in second-level analysis in SPM: a one sample t-test for I>0 (H0: I=0) and S>0 (H0:S=0) and a paired sample t-test comparing I and S (H0:I=S) was performed at every voxel. Using a voxel-wise false discovery correction for multiple comparison with q<0.05 and cluster-size threshold set at k>20 voxels, t-values greater than 2.81 and 2.77 respectively for I>0 and S>0 and 3.87 for I vs. S are significant.

### EEG data acquisition

The EEG was acquired using the actiCHamp (Brain products Gmbh Brain Products GmbH, Gilching, Germany) amplifier system with active electrodes. We recorded from a 128-electrode active array embedded in an elastic cap (ActiCap International, Inc.) in accordance with the 10–20 International System (AEEGS, 1991). In addition to the scalp electrodes, an active electrode was placed on the forehead (AFz, 25 mm above the nasion), and two electrodes on the left and right infraorbital rim to detect and clean eye movement artifacts. Impedance of all electrodes was kept below 5kΩ. The EEG signal was digitized at a sampling rate of 500 Hz (16 bit AD converter), and a hardware high-pass filtered was applied at 0.15 Hz to remove slow drifts.

Stimuli presented to the participants was identical to the one used for the fMRI study. The stimuli were presented on a High Definition screen kept approximately 1 meter away from the participants who were seated on a chair. Triggers were recorded at every camera change (corresponding to the transition between acts) during the movie. This was done by adding a white square on the last and first frame of each act on the side of the movie. An LED was then placed over the location of the square, and a wooden black masking frame ensured that the square was invisible to the participant. The output from the LED was then introduced as a digital channel input in the EEG recording system.

### EEG Data Analysis

EEG data was analyzed and preprocessed using in-house MATLAB (www.mathworks.com) routines and the FieldTrip analysis software in MATLAB (Oostenveld et al., 2011). The eye-movement artifact was removed using the Independent Component Analysis (ICA) procedure; a spatio-temporal ICA furnishes several topographical plots of the ICA components, which are then manually selected for eye-movement artifacts. These components are removed and the remaining components are back-projected to the original space to obtain the “cleaned” EEG signal. Since, the task did not involve any movement, we found almost no muscle artifacts in our data. Notch filters of 1 Hz centered on 50 Hz and its harmonics were used to remove the generic electrical cycle frequencies.

The Event-related potentials (ERP) were calculated around each camera change marking the beginning of a new act. Each trial was defined over a window 500 ms before and 1000 ms after the camera change. We therefore had 1292 trials in total including both the scrambled and the intact movies.

All trials were then averaged within condition. Because the movies are continuous, the 500 ms prior to a camera change is not a traditional baseline, but the end of the previous act. Regarding the period after the camera change, 98% of the acts had durations longer than 650ms. Accordingly, we focus our analyses on the first 650ms of this epoch, that is largely unperturbed by camera changes of subsequent acts.

To determine the time points on every channel for which the ERPs from the two conditions differed significantly, while addressing multiple comparison issues (750 time points × 128 channels), we used a standard cluster-based maxsum permutation test (Maris and Oostenveld, 2007 for an elaboration on this topic) as implemented in FieldTrip. For every channel and time-point the experimental conditions were compared by means of a t-test (with n=number of participants). All samples (channel-time pair) whose t-value are below a t-threshold corresponding to the uncorrected p<0.05 cluster-forming threshold are set to a value of zero. For all pairs exceeding this threshold, the value is first set to be the sum of its own t-value and those of the neighboring channels, integrated over a time period extending from −25ms to +25ms. Then in a second step, the value is set to the maximum of the sum values across these same spatio-temporal neighbors.

Once the max-sum statistic has been calculated we need to determine its likelihood under the null hypothesis using a Monte Carlo method. Trials of the different experimental conditions (intact and scrambled) are collected in a single set, as many trials from this combined data set as there were subjects in condition 1 are randomly drawn and placed into “pseudo-subset 1”. The remaining trials are placed in pseudo-subset 2. The test statistic (maxsum) is calculated for this random partition. The procedure for random partition and test statistics is repeated 5000 times and a histogram of the test statistic is constructed. We calculate the proportion of random partitions that resulted in a larger test statistic than the observed one. This proportion is the Monte Carlo significance probability, which is also our p-value. If this p-value for a particular time-channel pair is smaller than 0.05 we conclude that the data in the two experimental conditions are significantly different at that channel and time.

For testing the significance of time-invariant parameters, the same cluster approach was used, except that the summax was only applied across neighboring electrodes (and not also time).

### EEG Source Reconstruction

To capture the distributed representation of the underlying neuronal activity that resulted in the sensor-level measurements of brain activity, we performed source reconstruction using the minimum-norm estimation (MNE) method (Dale et al., 2000). MNE is an approach favored for evoked responses and for tracking widespread activity over time. It involves solving a distributed inverse solution that discretized the source space into locations in the brain volume using many current dipoles. It then estimates the amplitude of all modelled sources simultaneously to recover a source distribution whilst minimizing the overall source energy.

To perform the source reconstruction, we used an MNI template to create two geometric objects; the volume conduction model and the source model. The volume conduction model determines the physics of the propagation of electrical activity through the head, which in turn depends on the conductivity of the various tissues between the source and sensor. The source space, which will be populated with current dipoles, is the cortical sheet which is extracted from the anatomical image using a combination of FreeSurfer (https://surfer.nmr.mgh.harvard.edu/) and MNE Suite (http://martinos.org/mne/stable/index.html). The volume conduction and source models are then used to determine the lead fields (from source to sensor space) using the OPENMEEG (http://openmeeg.github.io/). The FieldTrip EEG analysis package was used to wrap the above packages along with helper functions to construct the pipeline for source reconstruction. The noise-covariance is calculated using a time-locked analysis over the sensor space. The leadfields along with the noise-covariance was used to reconstruct the source-level activity at every time-step.The ERPs of the two conditions (intact and scrambled) were contrasted to determine the time instances during which they differed significantly. The source reconstruction was performed at these time-points to reveal the source of the brain activity on the cortex.

## Acknowledgments

This work was supported by the Netherlands Organization for Scientific Research (VENI: 451-09-006, VIDI: 452-14-015 to V.G.), the Brain and Behavior Research Foundation (NARSAD young investigator 22453 to V.G.) and the European Research Council of the European Commission (ERC-StG-312511 to C.K.). We thank Judith Suttrup for acquiring and analyzing the data necessary to define the mirror network mask, Henk Stoffels and Abelraham Abdelgabar for helping with the stimulus recording, Juha Pajula for help with using the ISC toolbox. Correspondence should be addressed to.

## Supplementary Material

### Supplementary Methods and Results

Supplementary Table S1. Regions corresponding to ISC Intact > baseline.

Supplementary Table S2. Regions corresponding to ISC Scrambled > baseline.

Supplementary Table S3. Regions corresponding to bGLM Intact > baseline.

Supplementary Table S4. Regions corresponding to bGLM Scrambled > baseline.

Supplementary Methods S1. Putative mirror neuron system localizer.

Supplementary Methods S2. Motor localizer.

Supplementary Figure S1. Correlation between ISC-ROIs.

**Supplementary Table S1.**
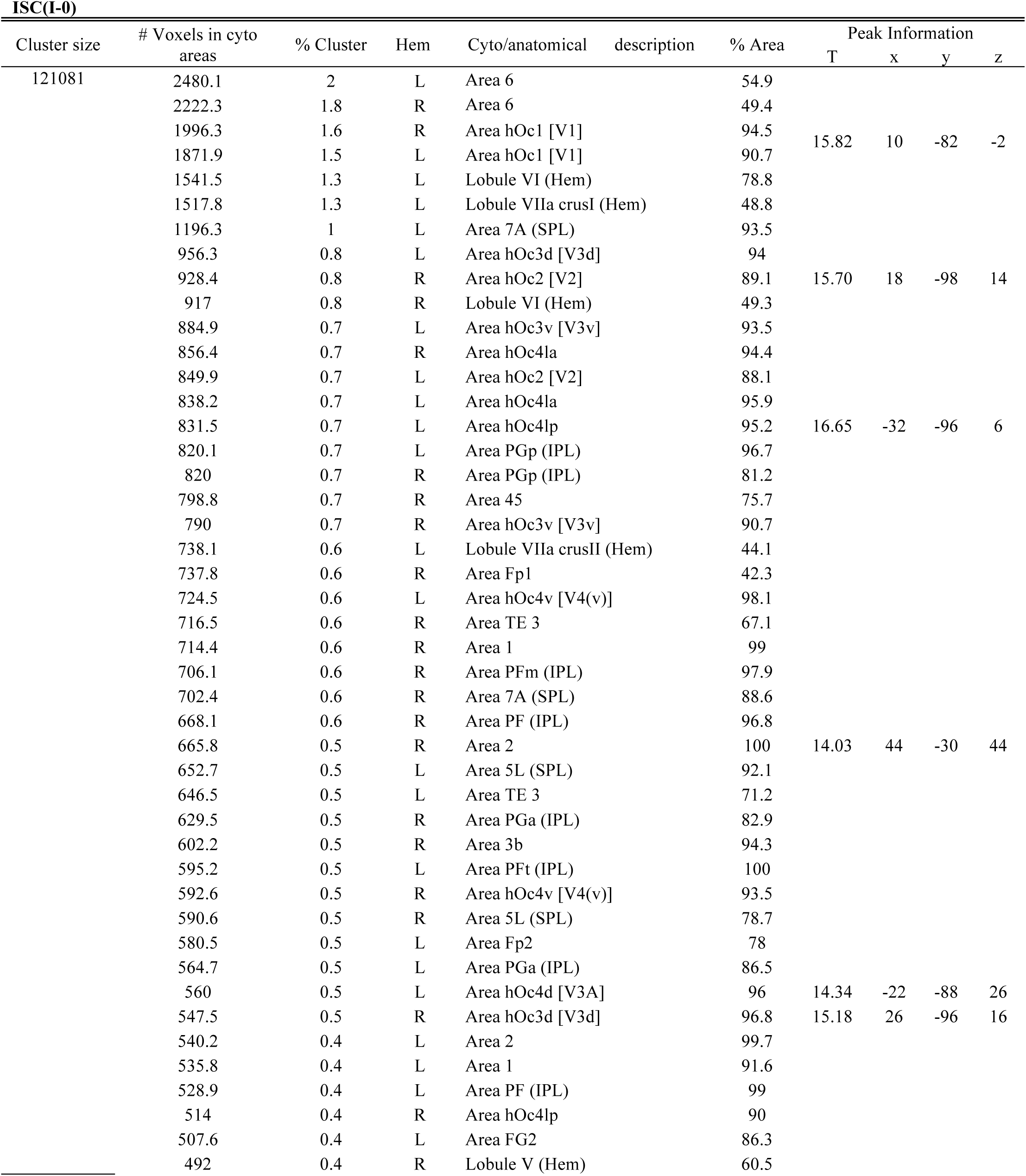
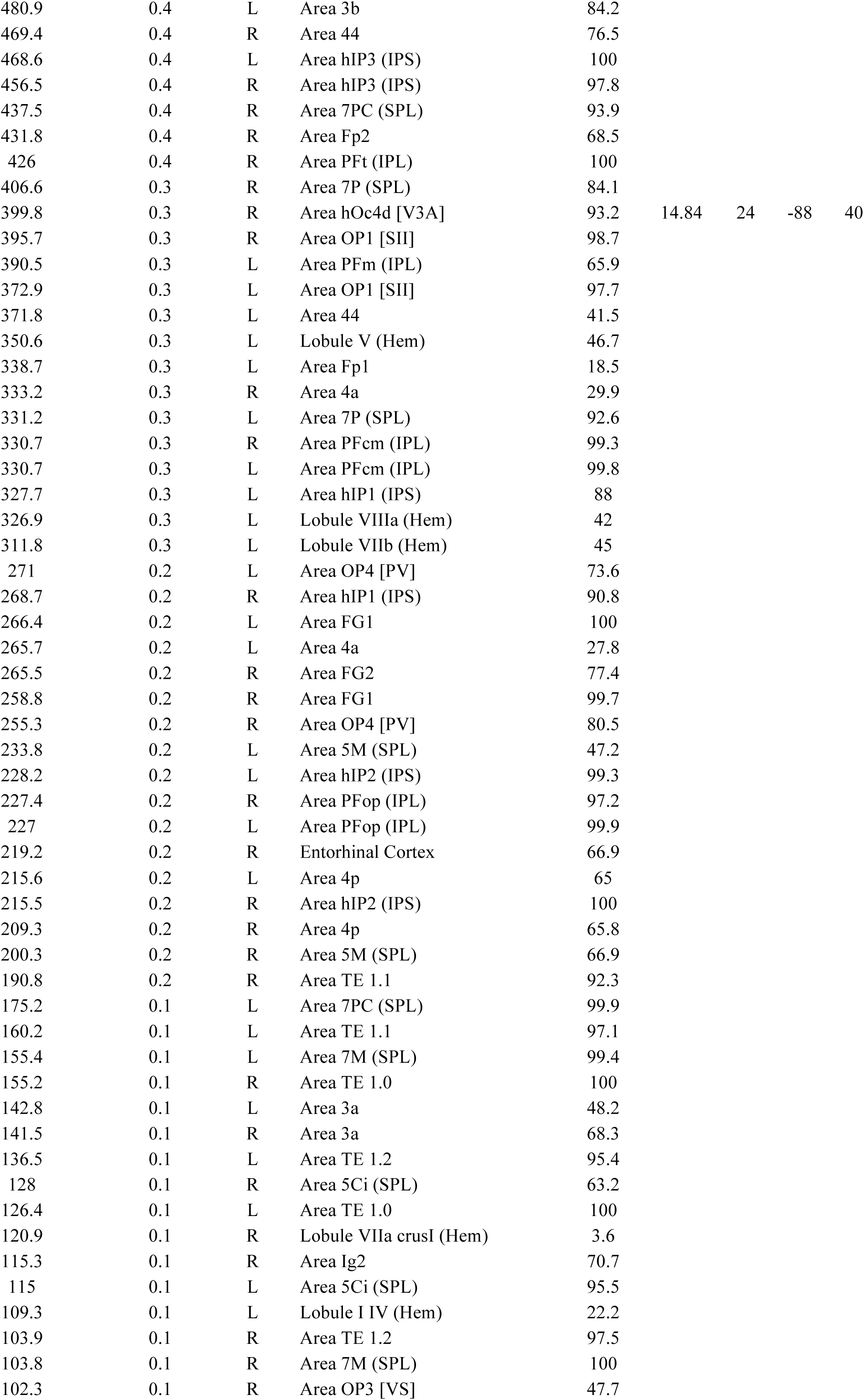
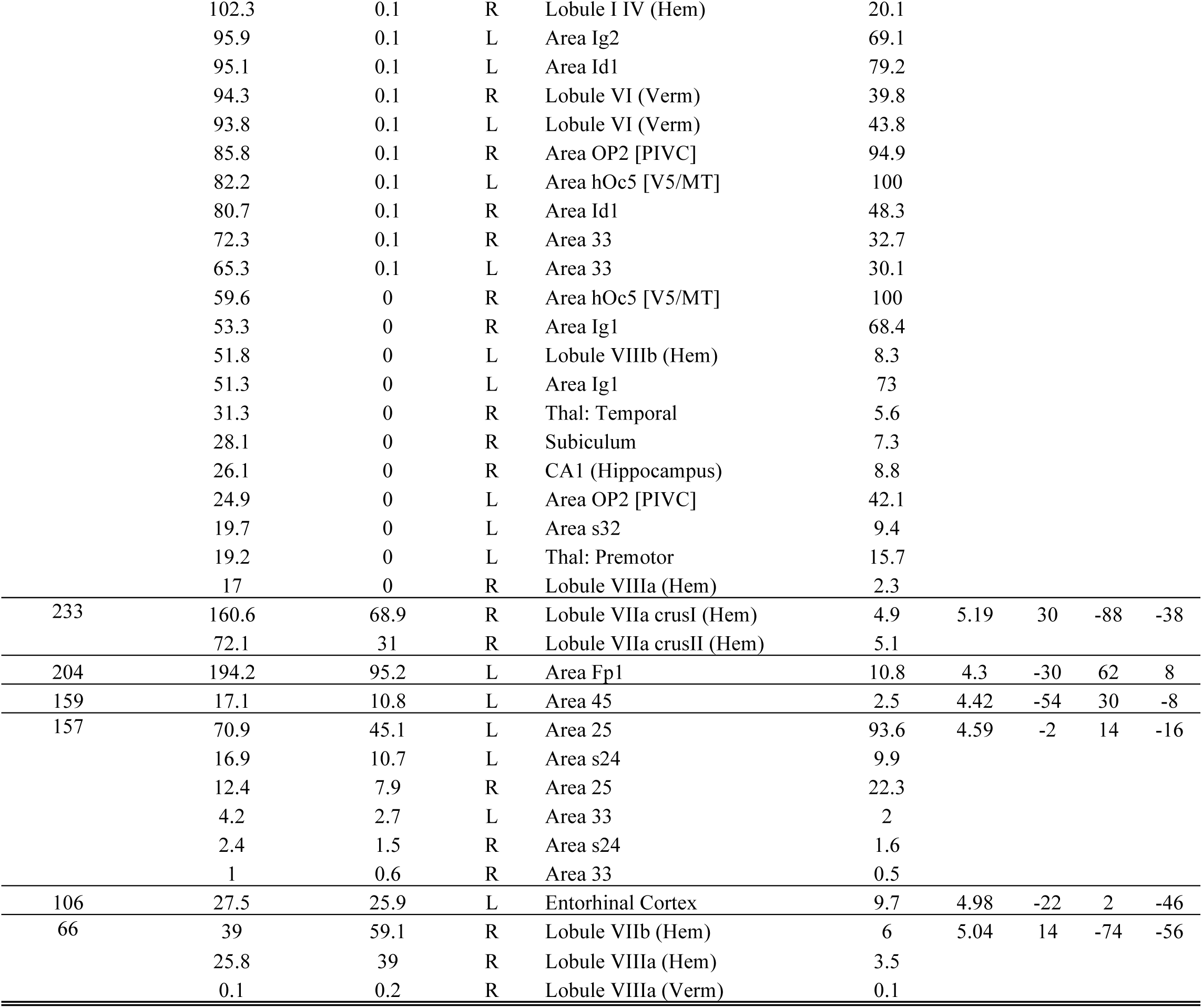
Regions with bGLM Intact > baseline labeled using SPM Anatomy Toolbox (P_uncr_<0.001, k>20 voxels, corresponding to t-values > 3.52; all voxels survived the q<0.05 threshold of t>2.77). Conventions as in Table 2 of the main manuscript.

**Supplementary Table S2.**
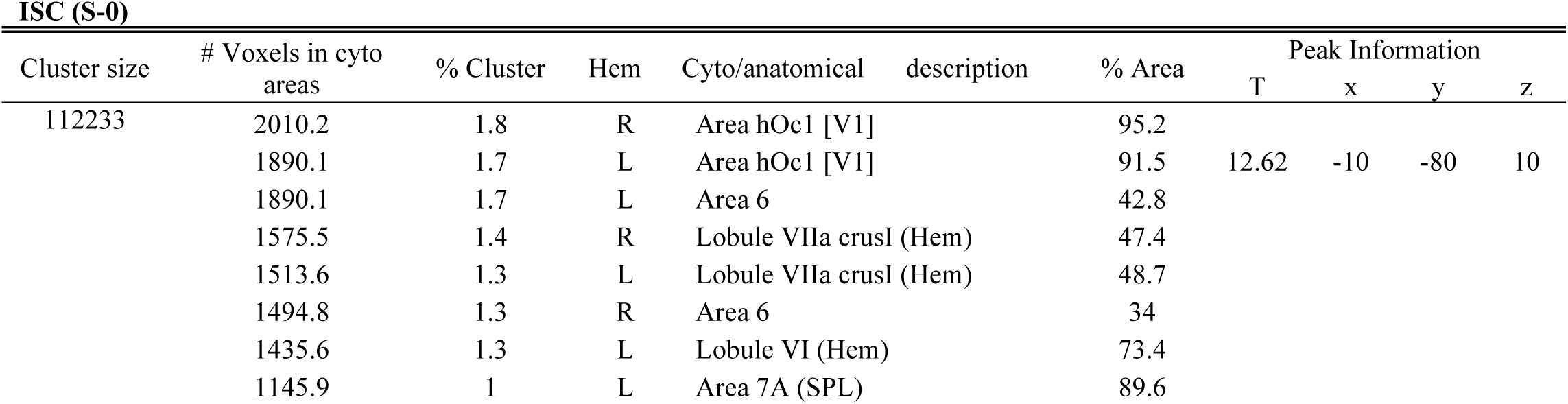
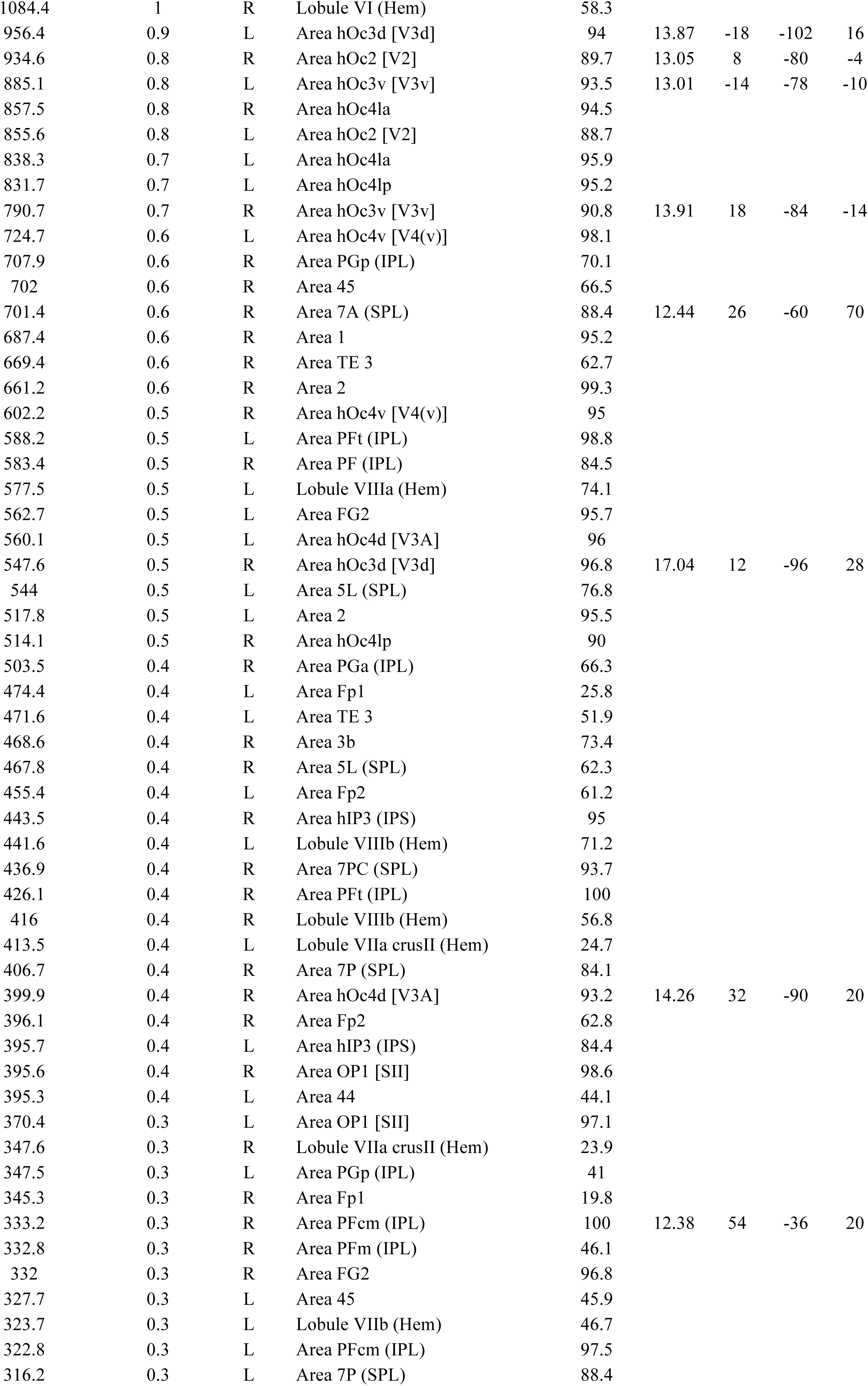
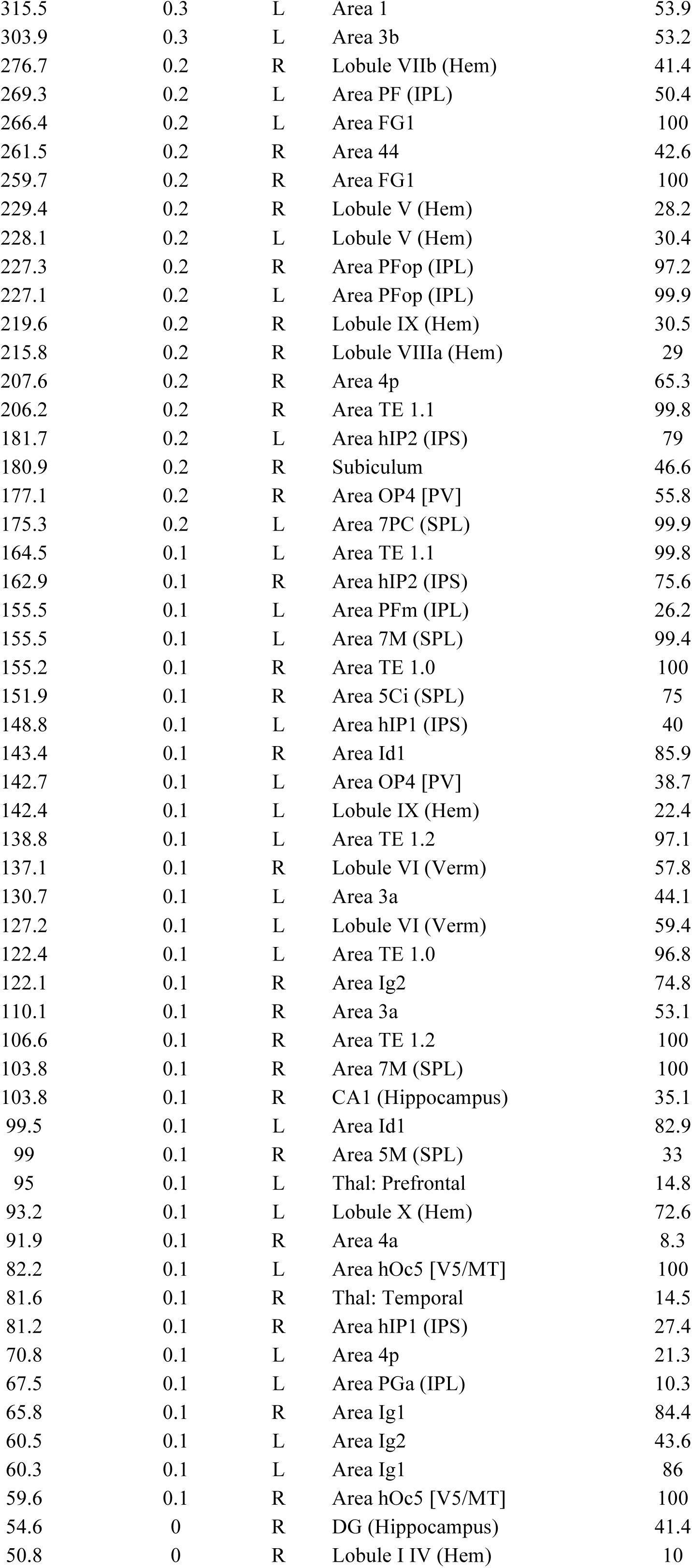
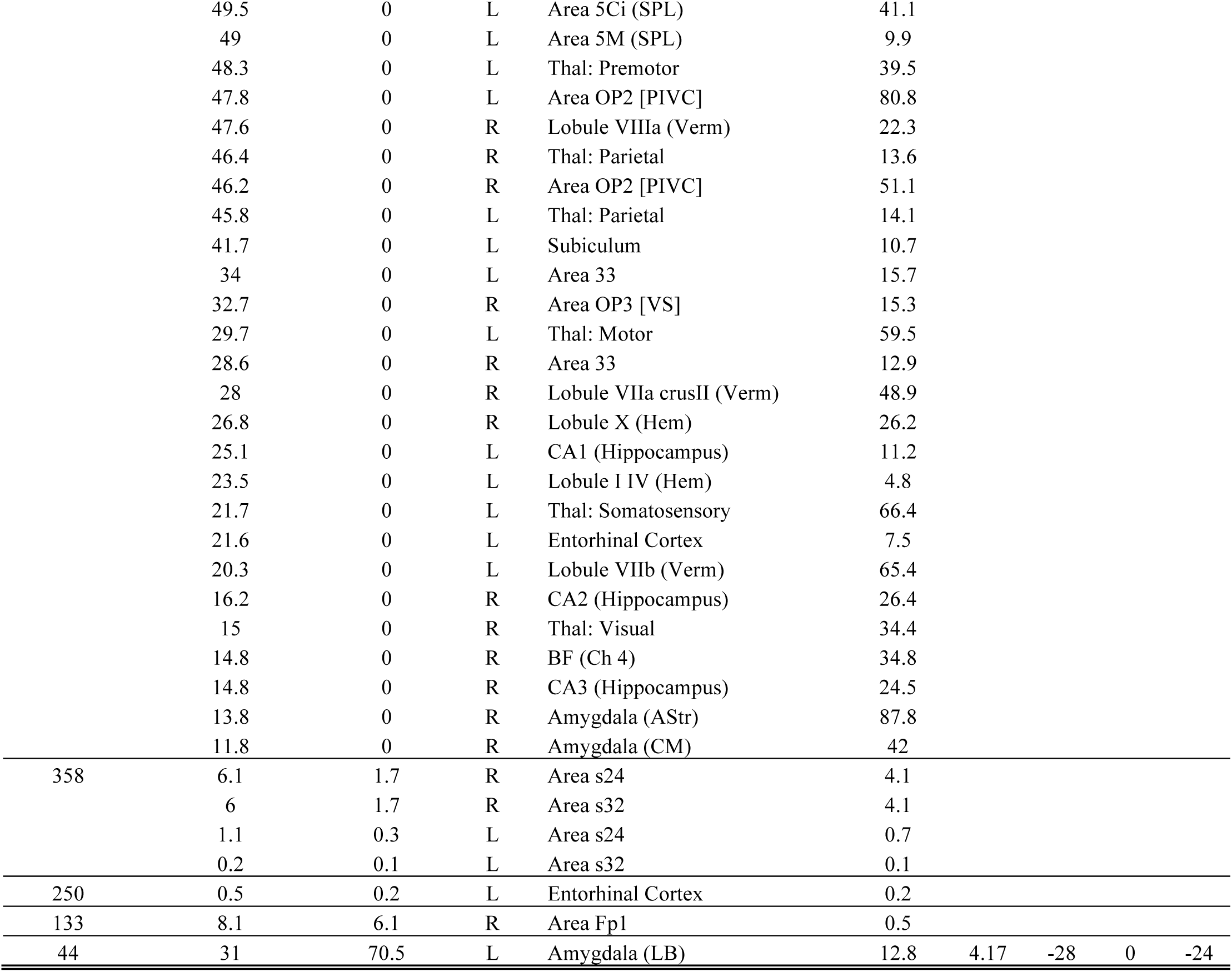
Regions with ISC Scrambled > baseline labeled using SPM Anatomy Toolbox (Puncr<0.001, k>20 voxels, corresponding to t-values > 3.52; all voxels survived the q<0.05 threshold of t>2.58). Conventions as in Table 2 of the main manuscript.

**Supplementary Table S3.** Regions with bGLM Intact > baseline labeled using SPM Anatomy Toolbox (Puncr<0.001, k>20 voxels, corresponding to t-values > 3.52; all voxels survived the q<0.05 threshold of t>2.81). Conventions as in Table 2 of the main manuscript.

| Cluster size | # Voxels in cyto areas | % Cluster | Hem | Cyto/anatomical description | % Area | Peak Information |  |  |  |
| --- | --- | --- | --- | --- | --- | --- | --- | --- | --- |
|  |  |  |  |  |  | T | x | y | z |
| 3251 | 382 | 11.8 | R | Area hOc4la | 43.1 | 9.11 | 38 | -80 | 4 |
|  | 371.4 | 11.4 | R | Area hOc4v [V4(v)] | 59.9 | 7.44 | 28 | -76 | -18 |
|  | 276.9 | 8.5 | R | Area hOc3v [V3v] | 32.5 | 16.11 | 26 | -90 | -10 |
|  | 193.8 | 6 | R | Area hOc4lp | 34.7 | 10.13 | 40 | -72 | -10 |
|  | 174.6 | 5.4 | R | Area FG1 | 68.8 | 5.15 | 32 | -62 | -14 |
|  | 124.2 | 3.8 | R | Area hOc1 [V1] | 6 |  |  |  |  |
|  | 119.9 | 3.7 | R | Area FG2 | 35.8 |  |  |  |  |
|  | 68.2 | 2.1 | R | Lobule VIIa crusI (Hem) | 2.1 |  |  |  |  |
|  | 61.2 | 1.9 | R | Area hOc2 [V2] | 6 |  |  |  |  |
|  | 48.6 | 1.5 | R | Area hOc5 [V5/MT] | 83.5 |  |  |  |  |
|  | 30.8 | 0.9 | R | Lobule VI (Hem) | 1.7 |  |  |  |  |
| 2532 | 429.4 | 17 | L | Area hOc4la | 50.2 | 10.05 | -46 | -74 | 0 |
|  | 319.1 | 12.6 | L | Area hOc4v [V4(v)] | 44.2 |  |  |  |  |
|  | 242.5 | 9.6 | L | Area hOc4lp | 28.4 | 6.23 | -30 | -90 | 12 |
|  | 205.1 | 8.1 | L | Area hOc3v [V3v] | 22.2 | 10.84 | -24 | -92 | -10 |
|  | 151 | 6 | L | Area FG2 | 26.3 |  |  |  |  |
|  | 103.5 | 4.1 | L | Area FG1 | 39.7 |  |  |  |  |
|  | 81.5 | 3.2 | L | Area hOc5 [V5/MT] | 101.4 |  |  |  |  |
|  | 31.4 | 1.2 | L | Lobule VIIa crusI (Hem) | 1 |  |  |  |  |
| 2272 | 504.2 | 22.2 | R | Area 2 | 77.4 | 7.28 | 38 | -34 | 40 |
|  | 290 | 12.8 | R | Area PFt (IPL) | 69.6 | 8.04 | 50 | -24 | 38 |
|  | 253 | 11.1 | R | Area 7PC (SPL) | 55.5 | 8.9 | 28 | -52 | 62 |
|  | 248.2 | 10.9 | R | Area 7A (SPL) | 32 |  |  |  |  |
|  | 198.5 | 8.7 | R | Area hIP3 (IPS) | 43.5 |  |  |  |  |
|  | 112.8 | 5 | R | Area 3b | 18.1 |  |  |  |  |
|  | 91.9 | 4 | R | Area PFop (IPL) | 40.2 |  |  |  |  |
|  | 88.6 | 3.9 | R | Area 1 | 12.6 |  |  |  |  |
|  | 45.9 | 2 | R | Area hIP2 (IPS) | 21.8 |  |  |  |  |
|  | 38.9 | 1.7 | R | Area 5L (SPL) | 5.3 |  |  |  |  |
|  | 36 | 1.6 | R | Area 3a | 17.8 |  |  |  |  |
|  | 10.4 | 0.5 | R | Area 7P (SPL) | 2.2 |  |  |  |  |
| 2013 | 457.2 | 22.7 | L | Area 7A (SPL) | 36.6 | 8.65 | -34 | -50 | 56 |
|  | 341.9 | 17 | L | Area PFt (IPL) | 58.7 | 7.07 | -44 | -32 | 38 |
|  | 231.4 | 11.5 | L | Area 2 | 43.7 | 7.08 | -38 | -42 | 52 |
|  | 175.8 | 8.7 | L | Area hIP3 (IPS) | 38.4 |  |  |  |  |
|  | 130.5 | 6.5 | L | Area 7PC (SPL) | 76.1 |  |  |  |  |
|  | 76.1 | 3.8 | L | Area 5L (SPL) | 11 |  |  |  |  |
|  | 29.4 | 1.5 | L | Area hIP1 (IPS) | 8.1 |  |  |  |  |
|  | 17.6 | 0.9 | L | Area PFop (IPL) | 7.9 |  |  |  |  |
|  | 15.1 | 0.8 | L | Area hIP2 (IPS) | 6.7 |  |  |  |  |
|  | 12.4 | 0.6 | L | Area 3b | 2.2 |  |  |  |  |
|  | 11.6 | 0.6 | L | Area 1 | 2 |  |  |  |  |
| 123 | 16.8 | 13.6 | L | Area 44 | 1.9 | 5.35 | -54 | 4 | 34 |
| 51 | 7.9 | 15.4 | R | Area 44 | 1.3 | 4.43 | 56 | 6 | 28 |
| 33 | 32 | 97 | L | Lobule VI (Hem) | 1.7 | 5.75 | -30 | -52 | -24 |
|  | 0.5 | 1.5 | L | Area FG2 | 0.1 |  |  |  |  |

**Supplementary Table S4.** Regions with bGLM Intact > baseline labeled using SPM Anatomy Toolbox (Puncr<0.001, k>20 voxels, corresponding to t-values > 3.52; all voxels survived the q<0.05 threshold of t>2.81). Conventions as in Table 2 of the main manuscript.

bGLM (S-0)
| Cluster size | # Voxels in cyto areas | % Cluster | Hem | Cyto/anatomical description | % Area | Peak Information |  |  |  |
| --- | --- | --- | --- | --- | --- | --- | --- | --- | --- |
|  |  |  |  |  |  | T | x | y | z |
| 5563 | 448.6 | 8.1 | L | Area hOc4la | 52.5 | 9.53 | -40 | -76 | -8 |
|  | 389.8 | 7 | L | Area hOc4v [V4(v)] | 54 |  |  |  |  |
|  | 364.9 | 6.6 | L | Area 7A (SPL) | 29.2 |  |  |  |  |
|  | 275.8 | 5 | L | Area hOc4lp | 32.3 | 8.45 | -30 | -90 | 6 |
|  | 252.1 | 4.5 | L | Area FG2 | 43.8 |  |  |  |  |
|  | 239.9 | 4.3 | L | Area hOc3v [V3v] | 25.9 | 12.37 | -22 | -88 | -8 |
|  | 239.1 | 4.3 | L | Area hIP3 (IPS) | 52.2 |  |  |  |  |
|  | 226.9 | 4.1 | L | Area PFt (IPL) | 39 |  |  |  |  |
|  | 204.5 | 3.7 | L | Area FG1 | 78.5 | 8.05 | -32 | -72 | -8 |
|  | 163.2 | 2.9 | L | Area 2 | 30.8 |  |  |  |  |
|  | 131.4 | 2.4 | L | Area 7PC (SPL) | 76.6 |  |  |  |  |
|  | 125.5 | 2.3 | L | Lobule VIIa crusI (Hem) | 4.1 |  |  |  |  |
|  | 104 | 1.9 | L | Lobule VI (Hem) | 5.4 |  |  |  |  |
|  | 82 | 1.5 | L | Area hOc5 [V5/MT] | 102 | 10.13 | -46 | -68 | 4 |
|  | 41.9 | 0.8 | L | Area hIP1 (IPS) | 11.5 |  |  |  |  |
|  | 36.8 | 0.7 | L | Area 5L (SPL) | 5.3 |  |  |  |  |
|  | 17.1 | 0.3 | L | Area hOc2 [V2] | 1.8 |  |  |  |  |
|  | 16.9 | 0.3 | L | Area hOc4d [V3A] | 3 |  |  |  |  |
|  | 16.4 | 0.3 | L | Area hOc1 [V1] | 0.8 |  |  |  |  |
|  | 12.9 | 0.2 | L | Area hIP2 (IPS) | 5.7 |  |  |  |  |
| 4050 | 409 | 10.1 | R | Area hOc4v [V4(v)] | 66 | 13.01 | 26 | -92 | -2 |
|  | 401.6 | 9.9 | R | Area hOc4la | 45.3 | 11.99 | 40 | -74 | -10 |
|  | 298.9 | 7.4 | R | Area hOc3v [V3v] | 35.1 | 14.14 | 24 | -88 | -10 |
|  | 237.5 | 5.9 | R | Area FG1 | 93.5 |  |  |  |  |
|  | 195.1 | 4.8 | R | Area hOc4lp | 34.9 |  |  |  |  |
|  | 181 | 4.5 | R | Area FG2 | 54 |  |  |  |  |
|  | 140.9 | 3.5 | R | Area hOc1 [V1] | 6.8 |  |  |  |  |
|  | 110.2 | 2.7 | R | Lobule VIIa crusI (Hem) | 3.4 |  |  |  |  |
|  | 69.2 | 1.7 | R | Area hOc2 [V2] | 6.8 |  |  |  |  |
|  | 56.8 | 1.4 | R | Lobule VI (Hem) | 3.1 |  |  |  |  |
|  | 48.6 | 1.2 | R | Area hOc5 [V5/MT] | 83.5 |  |  |  |  |
|  | 10.4 | 0.3 | R | Area hOc4d [V3A] | 2.5 |  |  |  |  |
| 2178 | 443.8 | 20.4 | R | Area 2 | 68.2 | 6.99 | 38 | -36 | 42 |
|  | 274 | 12.6 | R | Area 7A (SPL) | 35.3 | 6.92 | 24 | -58 | 50 |
|  | 259 | 11.9 | R | Area hIP3 (IPS) | 56.7 |  |  |  |  |
|  | 235.5 | 10.8 | R | Area PFt (IPL) | 56.5 |  |  |  |  |
|  | 225.9 | 10.4 | R | Area 7PC (SPL) | 49.5 | 8.21 | 30 | -52 | 60 |
|  | 105 | 4.8 | R | Area 3b | 16.8 |  |  |  |  |
|  | 52.6 | 2.4 | R | Area hIP2 (IPS) | 25 |  |  |  |  |
|  | 49.1 | 2.3 | R | Area 1 | 7 |  |  |  |  |
|  | 36 | 1.7 | R | Area 3a | 17.8 |  |  |  |  |
|  | 34.2 | 1.6 | R | Area PFop (IPL) | 15 | 8.29 | 50 | -26 | 38 |
|  | 24.2 | 1.1 | R | Area 7P (SPL) | 5.1 |  |  |  |  |
|  | 13 | 0.6 | R | Area 5L (SPL) | 1.8 |  |  |  |  |
| 62 | 1 | 1.6 | R | Area 44 | 0.2 | 4.77 | 52 | 6 | 30 |
| 24 | 9.2 | 38.5 | L | Area 44 | 1.1 | 3.99 | -50 | 4 | 30 |

## Supplementary Methods S1. Putative mirror neuron system localizer

The pMNS mask used in the current manuscript was generated from data collected on an independent sample of 35 participants who underwent an action execution and action observation task in MRI. The stimuli and tasks used with this sample of participants were identical to the Observation, Manipulate and Eye Execution conditions used by Arnstein et al. (Arnstein et al. 2011). The full study will be published separatly, but this data was used because it represents the largest sample ever used to localize the pMNS.

All participants signed an informed consent in accordance to the declaration of Helsinki, prior to the experiment, and met MRI safety requirements, had a normal or corrected to normal vision and had no history of neurological conditions or treatments. Four participants were excluded from the analysis: two because of excessive head motions, one reported sleepiness, and one because of image distortion. The remaining group was therefore composed of 31 participants.

### Observation task

Participants watched 39 unique movies of a human hand interacting with objects (ActionOBS), which were previously placed on a table (e.g. pouring wine into a glass). The 39 control movies displayed the same objects but the hand moved over the surface of the table without any meaningful object interaction (CtrlOBS). Each of the 7-sec blocks contained a series of three actions from the same condition. There were 13 blocks for each condition, which were separated by a centered fixation cross with a random duration of 8 to 12 seconds. The order of blocks was randomized across participants. The observation task was always acquired before the action execution task to avoid biasing action observation towards motor execution regions.

### Action execution task

Participants performed three of the actions presented in the observation task: stirring a spoon in a bowl, lifting a coffee cup as if to drink, and swirling a wine glass (ActionEXE). The plastic objects were placed on a t-shaped table secured above the waist of the participants. During scanning, participants performed actions in 13 blocks, with each block including the execution (in a pseudo-randomized order) of all three actions. Each action started and ended with the dominant hand being placed in a resting position at the edge of the t-shaped table. The appearance of a green circle displayed on a grey t-shaped background indicated the time at which to initiate the action. The circle position indicated which object to interact with (and therefore selected which of the three action to perform), and its duration paced the duration of the manipulation. When the circle turned red participants had to follow the motion of the circle with their eyes only (CtrlEXE).

For each participant, the MRI session included one anatomical scan (structural 3D spoiled gradient image of 170 slices; scan resolution = 256 × 256; field of view = 232 mm; voxel size: 1 × 1 × 1 mm) and two functional runs of 345 EPI volumes (one run for the observation, and one for the execution task) acquired in a Philips Intera 3T scanner, using a 32-channel coil. Functional images were acquired using an echo planar T2*-weighted gradient sequence covering 41 sequential axial slices (echo time = 28 ms; slice thickness = 3.5 mm; flip angle = 70°; repetition time = 2000 ms; scan resolution = 64 × 62; field of view = 224 mm; voxel size: 3.5 × 3.5 × 3.5 mm).

All analyses were run in SPM8 (Wellcome Trust Centre for Neuroimaging, UCL, UK) using Matlab 7.14 (The MathWorks Inc., Natick, USA). Data preprocessing included slice time correction, realignment and coregistration of functional and anatomical images to the mean functional image, normalization to MNI template through normalization parameters obtained by the segmentation of the anatomical image, and a 6×6×6 FMHW Gaussian kernel smoothing.

Separate general linear models were computed on the observation and execution data, each containing two boxcar predictors modeling the experimental conditions (ActionOBS, CtrlOBS or ActionEXE, CtrlEXE) and six predictors of no interest including the head movements determined during the realignment. The contrasts ActionOBS-CtrlOBS and ActionEXE-CtrlEXE were then computed at the subject-level. At the second, group level, we then performed a conjunction (of the conjunction-conjunction type) between two t-tests (ActionOBS-CtrlOBS>0, and ActionEXE-CtrlEXE>0) using a voxel-wise fdr correction that set a threshold of minimum t > 3.32. Only cluster of at least 20 voxels were retained in the pMNS mask.

## Supplementary Methods S2. Motor localizer

### Data acquisition and task for motor execution

Participants were the same as in the main fMRI experiment and data was acquired using the same fMRI acquisition methods on the same day after the main experiment. The participants were given toys of scale 3 cm to be held in both left and right hands comfortably during the scan. Instructions appeared as green crosses either on the left or right half of the screen signaling the participants to jostle the toy only in the respective hand of the cross for the duration of the appearance of the cross which was either two or three seconds. The inter-stimulus interval was randomly selected between 8 and 12 seconds. A total of 12 left and 12 right hand instructions were given during the scan. The entire task took 6 minutes to perform. In past experiment we found this simple localizer to robustly activate the motor and somatosensory system, but because of time constrains we were unable to include control conditions that would disentangle attentional and executive control related activations from motor processes proper, and results should thus be interpreted in the light of this lack of specificity. Preprocessing was performed as in the GLM analysis of the main experiment.

### Statistical analysis for motor localizer task

To investigate which of the ISC ROIs were significantly activated while participants themselves manipulated observed, after determining the 10 ROIs of interest for the ISC (I-S) condition as described in the main text, we extracted the Eigen time course of these ROIs using Marsbar. These time courses were placed into a GLM design with motion parameters (determined during realignment of the EPI volume) introduced as confounds. The motor execution was modelled as a boxcar regressor convolved with the canonical haemodynamic response function. Beta coefficients, reflecting the co-activation of the ROIs to the motor execution task, were tested for being different from zero. All ROIs that survived an FDR corrected t-value are highlighted with an asterisk against the ROI in figure 3 of the main text. Both the preprocessing and the statistical analysis was carried out using SPM12, Marsbar and in-house MATLAB routines.

## Supplementary Figure

**Figure S1.**
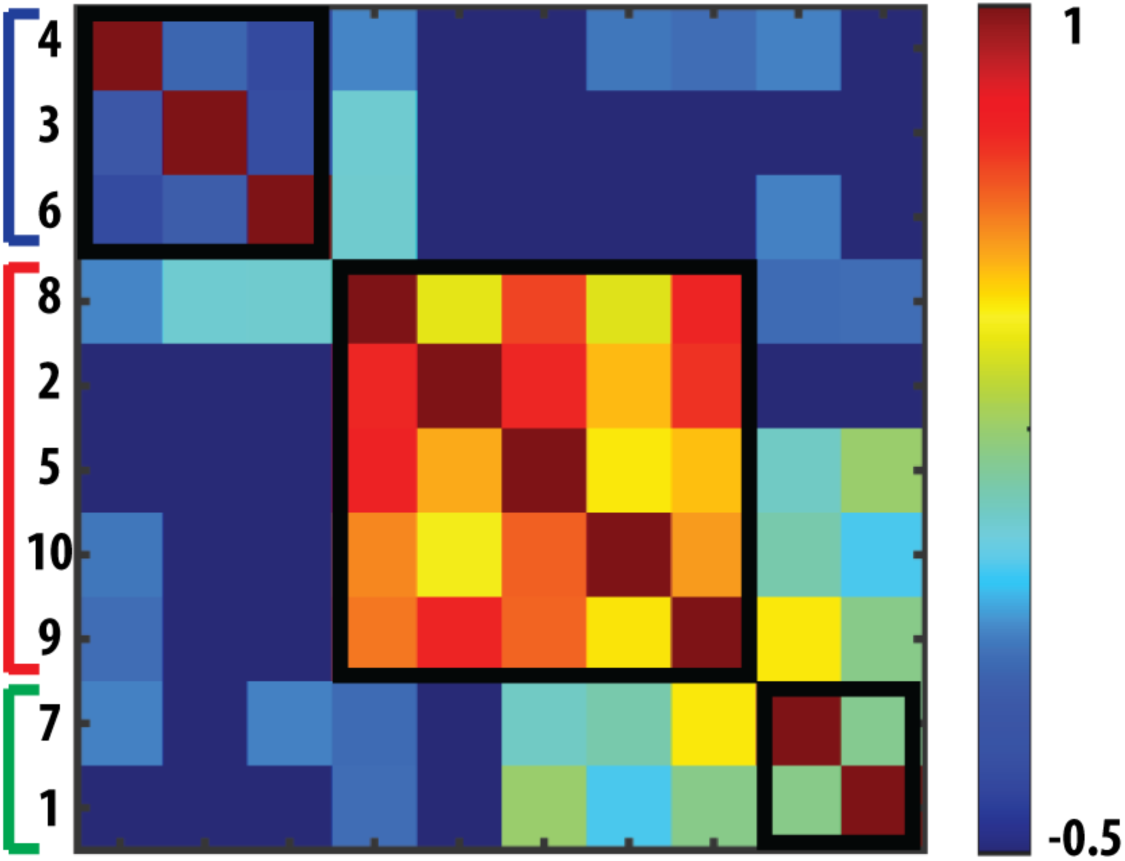
Correlation between ISC-ROIs. The Eigen timecourse of the each ROI (Figure 4) was extracted using Marsbar and the cross-correlation between them was calculated. The matrix of the correlation is plotted after spectral reordering. The ROI numbers correspond to Figure. 4 and are shown next to the rows. We can qualitatively see the three different clusters of ROIs. These were quantitatively established using the multi-dimensional scaling and k-means clustering procedure described in the main text, and evidenced by the black outline in the cross-correlation matrix.

